# Epidemic Risks associated with high-temperature replicating infectious hematopoietic necrosis virus (IHNV) field isolates: from molecular characterization to host pathogen interactions

**DOI:** 10.64898/2026.10.29.762392

**Authors:** Elise Gourhannic, Calvin Fauvet, Manon Mehraz, Pauline Pondaven, Mélissa Aurousseau, Maxence Fretaud, Stéphane Biacchesi, Jean K. Millet, Lénaïg Louboutin, Guillaume Lannuzel, Yannick Blanchard, Mai Lan Tran, Patrick Fach, Dimitri Rigaudeau, Thierry Morin, Christelle Langevin

## Abstract

Infectious hematopoietic necrosis virus (IHNV) is a WOAH-notifiable pathogen responsible for severe acute disease in salmonids at temperatures ranging from 4 to 14°C. At higher water temperatures, the disease is generally associated with reduced mortality and lower viral replication. In the context of climate change, rising water temperatures could favor the emergence of thermotolerant strains while representing a major physiological and metabolic stress potentially altering fish’s immune responses. This study aims to characterize the impact of increasing water temperatures on IHNV replication and host-pathogen interactions. Following pre-screening, the replication kinetics of the reference strain N61 and two field isolates RtNag0a and SA-22 were characterized in EPC (*Epithelioma Papulosum Cyprini*) and RTG-2 (Rainbow trout gonad) cell lines at 14, 20 and 22°C. Virulence and host-pathogen interactions were subsequently investigated *in vivo* in zebrafish (Zf) larvae using complementary microinjection and bath immersion infection models at 21.5°C and 23.5°C. *In vitro* studies revealed distinct thermal replication profiles, distinguishing thermosensitive from thermotolerant isolates, although these phenotypes were influenced by the cellular environment. The zebrafish model enabled the characterization of virulence and antiviral host responses associated with thermotolerant IHNV isolates. Notably, waterborne infection proved effective, contrasting with previous reports describing this exposure route as unsuitable for IHNV studies. These findings expand the growing list of emerging European IHNV genotypes capable of replicating at elevated temperatures (>20°C) and establish zebrafish larvae as a relevant model for virulence assessment of field isolates. Together, these results provide valuable insights into the potential impact of climate warming on IHNV epidemiology and support efforts to predict and anticipate future outbreaks under changing environmental conditions.

**Highlights:**

- Identification of thermotolerant IHNV isolates
- Zebrafish larvae recapitulate IHNV infection disease modelling host pathogen interactions
- *In vitro* replication profiles at high temperature and virulence in zebrafish are correlated
- Productive infection of zebrafish by immersion with clinical signs and mortality was obtained for the first-time using field IHNV isolates without prior *in vitro* thermal adaptation

## 1. Introduction

Aquaculture is one of the fastest-growing animal production sectors, yet its development and sustainability are challenged by diverse risks such as climate change, ecological crisis, and disease outbreaks. Meta-analysis conducted by Combe et al established the link between global warming and severity of viral infections in farmed aquatic animals (Combe et al., 2023). Rising temperatures alter ecosystem stability and promote pathogen emergence thereby threatening animal health and welfare. This underscores the need to re-evaluate the dynamics of host–pathogen–environment relationships for guiding the evolution of farming practices and for adapting therapeutic and preventive strategies to ensure resilience and sustainability in aquaculture systems. Among the viral fish diseases that constitute a significant concern, Infectious Hematopoietic Necrosis (IHN) is a severe infectious disease affecting both wild and farmed salmonid stocks, particularly impacting fry and juvenile stages but also adults to a minor extent (WOAH, 2026). In view of its significant impacts on animal health but also the economy and the environment, IHN has been designated as a notifiable disease by the World Organization for Animal Health (WOAH) and is listed in the EU Animal Health Law (Regulation EU 2016/429), thereby requiring the implementation of surveillance, control, and eradication measures in affected areas particularly in absence of vaccines suitable for juveniles and large-scale production systems in Europe. Over the past years, surveillance data in Europe shows an increase in IHNV incidence in trout farms and an expansion to previously unaffected Northern European countries (Abaddi et al., 2021; Kantala et al., 2026).

The transmission and dissemination of the disease are enhanced in intensive farming conditions, where fish are exposed to contaminated water and in close contact with infected individuals (horizontal transmission). The causative agent IHNV (ICTV taxonomic name: *Novirhabdovirus salmonid*) is a single stranded RNA virus (ICTV/International Committee on Taxonomy of Viruses 2025), belonging to the *Mononegavirales* order, in the *Rhabdoviridae* family and *Novirhabdovirus* genus. Its non-segmented genome of 11,200 nucleotides (nt) consists of six genes encoding five structural proteins: nucleoprotein (N), phosphoprotein (P), matrix protein (M), glycoprotein (G), RNA-dependent RNA polymerase (L) and a non-structural one, the non-virion protein (NV) (Kurath et al., 1985). At early stages of infection, the virus can be detected in fins, gills, skin, digestive system and blood before spreading to the hematopoietic organs (kidney, spleen, thymus, liver), the muscle and cartilage (Yamamoto & Clermont 1990; Chilmonczyk & Winton, 1994; Harmache, 2006; Dixon et al., 2016). Fins constitute highly permissive entry sites during waterborne exposure (Harmache et al., 2006), reinforcing the central role of superficial epithelial tissues in early IHNV infection. The clinical expression is characterized by lethargy, anorexia, skin melanosis and exophthalmos.

Based on glycoprotein (G) gene sequences, IHNV isolates are classified into five major genogroups: U (Upper), M (Middle), L (Lower) in North America, J (Japan), and E (Europe), reflecting their historical geographical distribution (Kurath et al., 2003; Bellec et al., 2017). The virus was first identified in North America, where U, M and L genogroups circulate among Pacific salmonid species, before spreading to Europe and Asia through international trade of infected eggs. Genogroup U has subsequently been reported in Japan, China and Russia (Rudakova et al., 2007; Huo et al., 2022), whereas M isolates have also been detected in China (Cieslak et al., 2017). Genogroup J comprises isolates recovered from rainbow trout in Japan and Korea (Nishizawa et al., 2006; Kim et al., 2007), and more recently in Kenya (Mulei et al., 2019). Noteworthy, phylogenetic analyses suggest that genogroup J evolved from genogroup U, while the European genogroup E originated from the M lineage following introduction into Europe (Enzmann et al., 2005). The prevalence of IHNV infection can reach high levels in endemic regions, with levels of virulence depending on the viral strain and genogroup, the environmental conditions (water temperature and fish density) as well as host-related factors including genetic background, developmental stage, microbiota and stress status.

IHN is considered to be a cold-water disease, with acute episodes occurring at low temperatures, typically between 4 and 14°C although evidence of replication at higher temperatures has been obtained under experimental challenges of fingerling rainbow trout or from epidemiological observations (Dixon et al., 2016; Vendramin et al., 2022). Global climate change is driving a rise in water temperatures creating a strong selective pressure that shapes RNA virus evolution through their inherently high mutation rates, as demonstrated both *ex vivo* and *in vivo* (Hoffmann et al., 2005; Mandary et al., 2019). The evolutionary plasticity of IHNV enables adaptation to temperature-dependent constraints, as evidenced by field isolates recovered at water temperature exceeding 14°C (Vendramin et al., 2022; Marsella et al., 2025). This thermal responsiveness facilitates the emergence and the selection of thermotolerant strains with modified fitness, transmissibility, and virulence (Troyer, 2002, Marsella et al., 2025).

Virulence evolution following host jump and host-associated temperatures (10°C and 15°C) was examined through a comparative analysis of 16 archival IHNV isolates representative of the U and M phylogenetic genogroups, respectively associated with sockeye salmon (*Oncorhynchus nerka*, ancestral host) and rainbow trout (*O. mykiss*, novel host) infections. The study showed that U genogroup infectivity remained higher in the ancestral host at 10°C, whereas M genogroup viruses adapted to the novel host and became more virulent than U at 15°C in rainbow trout (Loeher et al., 2025). More recently, Marsella et al. investigated the fitness and virulence features of six European isolates (E genogroup) selected based on their geographical origin, year of isolation, pathogenicity and water temperature, comprising a Danish variant responsible for recent severe outbreaks in Northern Europe (Vendramin et al., 2022) and Italian isolates from the library of Istituto Zooprofilattico Sperimentale delle Venezie (IZSVe) (Marsella et al., 2025). *In vitro* assays conducted across a temperature gradient from 15 to 25°C revealed distinct phenotypes from temperature-sensitive to thermotolerant isolates. The Italian IHNV isolate 67-1 and the Danish IHNV isolate DK-FF1/21–4070–1, collected in environments with temperatures of 16°C and 19°C, respectively, replicated at 15, 20, and 22°C, but not at 25°C. *In vivo* examination of these thermotolerant isolates was performed by challenging juvenile rainbow trout maintained at 12°C and 19°C following a two-week acclimation period. The infection caused by the Italian strain resulted in comparable mortality rates at both temperatures (around 90%), but with faster kinetics at 19°C. In contrast, the virulence of the Danish strain was attenuated at the highest temperature, yielding a survival rate of approximately 68 % at 19°C reduced to about 5 % at 12 °C; even though this strain replicated efficiently at 22°C *in vitro*. In parallel, analysis of the IHNV 224–2 isolate selected as a highly pathogenic cold water adapted IHNV revealed contrasting temperature-sensitivity phenotypes between *in vitro* and *in vivo* conditions. IHNV 224–2 replicated efficiently at 20 °C in cell culture but its virulence was substantially attenuated *in vivo* at 19°C, resulting in 78.8 % survival compared with only 14.3 % survival at 12°C. These results underscore the complexity of predicting temperature-dependent virulence of IHNV field isolates and highlight the limitations of extrapolating *in vitro* replication profiles to pathogenic outcomes in fish host.

Examination of IHNV isolates capable of replicating at increasing temperatures in the context of global changes demands the development of predictive experimental frameworks positioned at the interface between *in vitro* modelling and animal experimentation. Establishing such integrative systems is now essential both for improving biological relevance and for reducing reliance on animal use. As such, zebrafish (Zf) larvae, considered as an alternative to animal experimentation (Strähle et al., 2012), have been exploited to model IHNV infections and investigate host-pathogen interactions. Its optical transparency, small body size and associated tractable genome-editing technologies provide major advantages for functional analyses (Aggad et al., 2009 & 2010; Ludwig et al., 2011; Langevin et al., 2013; Briolat et al., 2014) and detailed characterization of the dynamic of infectious processes and immune responses (Ludwig et al., 2011). Its tolerance to a wide temperature range is of particular interest by model infection to a broad diversity of viruses (Sullivan et al., 2021), reinforcing Zf larvae as a powerful system for dissecting host–virus interactions across diverse thermal niches (from 20 to 33°C). IHNV Zf infection models have been successfully established at 20°C for juveniles and adults using both bath and injection administration modes (La Patra et al., 2000) whereas 24°C has been employed for effective infection of embryonic stages following intravenous injection (Aggad et al., 2009). Under these conditions, a Canadian IHNV (IHNV 25-70) strain progressively adapted *in vitro* to 25°C enabled detailed characterization of virus-induced interferon signaling pathways as well as comprehensive *in vivo* analyses of viral replication and dissemination together with the associated clinical signs typically elicited during disease progression (Aggad et al., 2009, Ludwig et al., 2011, Briolat V., 2014). Temperature shift experiments from 24°C to non-permissive 28°C demonstrated the impact of thermal regime on disease progression as a 4°C increase was sufficient to prevent viral spreading and interfered with the viral life cycle as well as host immune responses (Ludwig et al., 2011).

In the present study, we investigated the impact of increasing water temperatures on IHNV replication, fitness, virulence and host defense mechanisms using complementary *in vitro* and *in vivo* approaches. Across six IHNV isolates originating from the WOAH and French reference laboratories, two isolates replicated in a permissive EPC *Epithelioma Papulosum Cyprini* cell line at 20 and 22°C and induced cytopathic effects. Their pathogenicity was then evaluated in Zf larvae at 21 and 23.5°C revealing disease severity consistent with *in vitro* replication profiles at both temperatures. Using reverse genetics, a recombinant virus encoding the mCherry fluorescent reporter protein was generated, enabling *in vivo* imaging of the infection process and visualization of viral entry, replication, and dissemination throughout zebrafish larvae. In addition, *Tg(mpx*:*GFP)* transgenic larvae expressing fluorescent neutrophils were used to monitor host responses during infection, allowing the visualization of neutrophil recruitment, distribution, and interactions within infected tissues. By expanding the list of European IHNV genotypes capable of replication at temperatures exceeding 20°C and validating zebrafish larvae as a model for virulence studies, this work advances our understanding of the epidemiological consequences of climate warming and supports the development of predictive approaches for future disease outbreaks.

## 2. Materials and Methods

### 2.1. Cells and IHNV field isolates

*Epithelioma Papulosum Cyprini* (EPC) and Rainbow Trout Gonad (RTG-2) cell lines were maintained at 20 and 14°C respectively in a Glasgow’s modified Eagle’s medium (MEM Glasgow powder (Pan Biotech) with 0.19 M Tris-HCl and HCO3Na (Sigma); pH 7.6) supplemented with 10% fetal bovine serum (Eurobio), 1X antibiotics (100 IU/mL penicillin G, 0.1 mg/mL streptomycin) and 10% l-glutamine (HyClone). These two cell lines are permissive to IHNV and have optimal thermal growth limits ranging from 15°C to 33°C and from 4°C to 26°C for EPC and RTG-2, respectively (Fijan et al., 1983; Wolf et al., 1962). Six IHNV isolates were initially selected: 1075b (*in vitro* passage P1), SA-22-05674 (P1) and N61 (P3) from genogroup E – this last one used as the French reference from genogroup E; FR0031 (P3) from genogroup L; WRAC (P3) from genogroup M; RtNag0a (P3) from genogroup J; the first three originated from the French Reference Laboratory for regulated fish diseases and the three last from the WOAH reference laboratory for IHNV (Western Fisheries Research Center, Seattle). Selection criteria incorporated geographical diversity, genotypes, species from which they were isolated and data concerning experimental virulence on rainbow trout (when available; Table 1). These strains were grown on an EPC cell monolayer at 14°C ± 2°C for 6–8 days. Once the cytopathic effect (CPE) was complete, cell culture supernatants were recovered, centrifuged 15 min at 2,000 g at 4°C ± 3 then stored at -65°C until use. An aliquot of each viral suspension was titrated using the 50% tissue culture infective dose (TCID_50_) endpoint method in 96-well plates, and calculated as described by Kärber (1931). The infectious titers of the viral production were found to be between 2.25 × 10^7^ and 1.26 × 10^9^ TCID_50_/mL.

**Table 1.** Selection of IHNV field isolates based on geographic and genetic diversities, host species and virulence levels reported on rainbow trout.

| Identification | Origin | Host species | Virus genotype | Year of isolation | T° of isolation (°C) | Cumulative mortality on RT (%) | Reference GenBank |
|---|---|---|---|---|---|---|---|
| N61 | France | RT | E | 1989 | nd | 47 | L40883.1 |
| SA-22-05674 | France | RT | E | 2022 | 14 | nd | SUB165469 |
| 1075b | France | RT | E | 2017 | 13 | nd | MN083251.1 |
| FR0031 | USA | Chinook salmon | L | 2000 | nd | 90 | X |
| WRAC | USA | RT | M | 1983 | nd | 58 | NC_001652.1 |
| RtNag0a | Japan | RT | J | 2006 | nd | 8 | X |

RT: rainbow trout. nd: undetermined.
X:  Sequence currently being deposited in GenBank

### 2.2. Phylogenetic analyses

A dataset comprising 47 viral genomes was used for the phylogenetic analysis, including the six IHNV isolates selected in this study together with representative IHNV reference sequences retrieved from GenBank and the published literature. The sequences were assembled and inspected using Geneious Prime (version 2026.x), subsequently aligned using the MAFFT multiple sequence alignment algorithm on the Galaxy platform and poorly aligned regions and gaps were finally removed using TrimAl (version 1.5.0). The quality of the final alignment was visually inspected prior to phylogenetic analysis. Phylogenetic relationships were inferred using the Maximum Likelihood (ML) method implemented in IQ-TREE 3.1.3 (edition built 19 June 2026) on the Galaxy platform with default parameters. Branch support was assessed using 1,000 bootstrap replicates, and only bootstrap values ≥ 70% were considered statistically supported. The resulting phylogenetic tree was exported in Newick format and visualized using iTOL (Interactive Tree Of Life), where it was rooted using the midpoint rooting method. The major clades were color-coded for clarity.

A comparative phylogenetic analysis was additionally performed including the IHNV isolates IHNV/O.mykiss/I/TN/534/Dec1 (abbreviated here IHNV 534), IHNV/O.mykiss/I/TN/224–2/Mar18 (IHNV 224-2), IT-217A (IHNV 217/A), IHNV DK-FF1/21–4070-1 (IHNV DK) and IHNV/O.mykiss/I/BG/67–1/Feb22 (IHNV 67-1) (Marsella et al., 2025); IHNV/143/14 (Abaddi et al., 2021); and the experimentally thermo-adapted IHNV25 variant recently described by Fauvet et al. (submitted). This analysis was conducted using the same approach as described above, based on complete glycoprotein (G) gene sequences. In addition, partial amino acid sequences of the viral polymerase (L) were aligned and compared using MEGA 11 software, except for isolate IHNV 143/14 for which the sequence is not available.

### 2.3. Construction of IHNV variant cDNA clones and recombinant virus recovery

A full-length cDNA copy of the RNA genome of the SA-22 isolate was synthesized *in vitro* and cloned in a pUC57 plasmid (Genscript). The resulting construct, referred to as pSA-22IHNV, consisted of a full-length antigenomic cDNA copy of the IHNV variant RNA genome cloned under the control of a T7 promoter, and downstream a hepatitis delta virus ribozyme sequence and a T7 transcription terminator, as previously reported (Biacchesi 2000). These constructs were subsequently modified to introduce an expression cassette encoding the fluorescent protein mCherry, as previously described (Biacchesi 2010). The insertion of a fluorescent reporter cassette into the viral genome allows direct visualization of infected cells and tissues, enabling real-time monitoring of viral replication and dissemination. In this study, the mCherry expression cassette was inserted into the intergenic region between the M and G genes of pSA-22IHNV, leading to the construct pSA-22IHNV-mCherry. This recombinant virus was designed to facilitate tracking of infection dynamics, analysis of viral spread, and characterization of host–virus interactions both *in vitro* and *in vivo*. Recombinant viruses were recovered from the full-length antigenomic constructs using a reverse genetics approach, as previously described (Biacchesi, 2000). Briefly, EPC cells were first infected with a recombinant vaccinia virus expressing the T7 RNA polymerase (vTF7-3, Fuerst et al., 1986, kindly provided by Dr B. Moss) and then co-transfected with the pSA-22IHNV-mCherry plasmid together with helper plasmids encoding the viral N, P, and L proteins under the control of the T7 promoter. The expression of these viral proteins, in combination with the antigenomic RNA transcribed from the full-length construct, enables the formation of functional ribonucleoprotein complexes and the initiation of viral replication. Following transfection, cells were incubated at 37°C for 6 h and then shifted under IHNV optimal temperature condition (15°C) and monitored for the appearance of fluorescence. Successful recovery of the recombinant virus was confirmed by live cell imaging upon detection of the mCherry fluorescence signal under UV illumination using a Carl Zeiss fluorescence microscope equipped with a 5X objective, indicating active viral replication and spread within the cell monolayer.

Cell culture supernatants were subsequently collected and used to amplify viral stocks through two successive passages on fresh EPC cells. Viral stocks obtained after two passages were then used for downstream experiments. The presence of the mCherry reporter and viral infectivity were verified based on fluorescence detection and propagation efficiency in cell culture.

### 2.4. In vitro infection at different temperatures

Viral replication was assessed on EPC at four temperatures: 14°C (optimal temperature), 20°C, 22°C, and 24°C (suboptimal temperatures) using metrologically verified incubators (COFRAC accreditation) equipped with temperature probes ensuring continuous monitoring (OCEAView system). Cells were seeded in 96-well plates at a density of 2 × 10^5^ cells per well. After 24 hours of incubation at 29°C, confluent monolayers were infected at the targeted temperature with each viral strain at a multiplicity of infection (MOI) of 0.1. At 1-, 2-, 3-, 6-, and 8-days post-infection (dpi), CPE was monitored and plates were subsequently frozen at −80°C to induce cell lysis. Supernatants corresponding to intracellular and secreted viral particles were then collected to determine infectious viral titers as described by Kärber (1931). All six IHNV isolates were first evaluated in a preliminary screening assay across the tested temperatures. Based on the observed phenotypic differences, three isolates (N61, SA-22, and RtNag0a) were subsequently selected for detailed characterization in triplicate. These experiments were reproduced in RTG-2 under the same experimental conditions at both 14°C and 20°C in triplicate.

Replication kinetics of N61, SA-22 and RtNag0a were examined at earlier time points in EPC cells in 6-well plates infected at an MOI of 0.1. The viral inoculum was adsorbed onto cell monolayer in a final volume of 1 mL for 1 h. Following inoculum removal, the monolayers were rinsed with PBS before addition of 3 mL culture medium. At 0-, 24-, 48- and 72-hours post-infection (hpi), supernatants containing excreted viral particles were collected before rinsing followed by lysis of the cell monolayer at −80°C to recover intracellular virus particles. Infectious viral titers were determined in supernatant fractions (Kärber, 1931). CPE was monitored by microscopy throughout the experiment. A total of 5 biological replicates were done per isolate.

### 2.5. In vivo experimentations

#### 2.5.1 Ethic statements

All animal experiments were conducted in strict compliance with European regulations and guidelines governing animal experimentation and welfare (European Union Directive 2010/63). Experimental protocols were reviewed and approved by the local ethics committee for animal experimentation (Ethics Committee for Animal Experimentation at the INRAE Research Center in Jouy-en-Josas; COMETHEA INRAE no. 45) and authorized by the French Ministry of National Education, Higher Education, and Research (authorization numbers: APAFIS#29801–2021021110262075 v2 and APAFIS#IERPZEBRA). Fish facilities were also approved under authorization number C78-720. To reduce animal suffering and distress, all procedures were performed under light anesthesia, achieved by bath immersion in tricaine (MS-222 at 0.025%, Sigma). Fish were daily monitored for clinical signs and survival. Animals exhibiting characteristic symptoms of infection were humanely euthanized by bath immersion in a lethal dose of tricaine (MS-222, Sigma; 0.125%).

#### 2.5.2 Zebrafish housing, husbandry and experimental challenges

Zebrafish (*Danio rerio;* Zf) were maintained at 28° C in freshwater under a 14 h light/10 h dark photoperiod. They were housed in the animal facilities designed and operated in accordance with applicable local animal welfare regulations (see above). The lines used in this study included nacre^w2/w2^ (Lister et al., 1999) and *Tg(mpx*:*GFP)^i114e^* (Renshaw et al., 2006).

At 3 days post-fertilization (dpf), Zf larvae were infected with IHNV isolates N61, SA-22 or RtNag0a. Larvae were anesthetized with tricaine (0.025 %) and positioned on a plate containing 1% Phytagel (Sigma-Aldrich) diluted in E3 medium (for Zf larvae, Cold Spring Harbor Protocols, 2011) to enable individual microinjection. Injections were carried out using a glass microneedle mounted on a mechanical micromanipulator (M-152, Narishige) and a FemtoJet microinjector (Eppendorf). Larvae were injected into the caudal vein with 2 nL of IHNV inoculum (10^7^ TCID_50_ per larvae) or 2 nL of cell culture medium (mock-infected larvae) up to 20 larvae per condition. Following injection, larvae were maintained in E3 medium at 21.5 or 23.5°C. Alternatively, a waterborne model of infection was developed by incubating 20 zebrafish larvae at 3 dpf in 3 µL of low-passage viral stocks diluted in a final volume of 100 µL of E3 medium for 1 h at 21.5 or 23.5°C, resulting in final virus concentrations of ∼1.07 × 10^7^ TCID_50_/mL, ∼3.78 × 10^7^ TCID_50_/mL and ∼2.13 × 10^6^ TCID_50_/mL for N61, SA-22 and RtNag0a, respectively. Mock infected larvae were exposed to cell culture media. Larvae were monitored daily to observe clinical signs and to record mortality rates. Each experiment was performed in triplicate.

#### 2.5.3 Determination of infectious titers by plaque assay

Infectious viral titers were determined by plaque assay at 0-, 6-, 24-, and 48-hpi as described by Souto et al (2024). Four larvae were collected at each time point, providing four biological samples per time point per experiment. Larvae were dried and stored at −80°C until analysis. Individual larvae were homogenized in 100 µL of cell culture medium supplemented with gentamicin (Sigma) using sterile ceramic beads and a Precellys 24 Touch tissue homogenizer (Bertin Technologies) for two cycles of 15 s at 6,000 rpm. Homogenates were clarified by centrifugation for 3 min at 10,000 rpm. The resulting lysates were 10-fold serially diluted in cell culture medium supplemented with 2% FBS and used to infect EPC cells seeded in 96-well plates. Viral titers were determined by plaque assay and expressed as plaque-forming units per milliliter (PFU/mL).

### 2.6. Transcriptomic analyses

The primers and probes used for reverse-transcription (RT) – quantitative polymerase chain reaction (qPCR) analyses conducted on *in vitro* and *in vivo* samples are listed in Table 2.

**Table 2.**
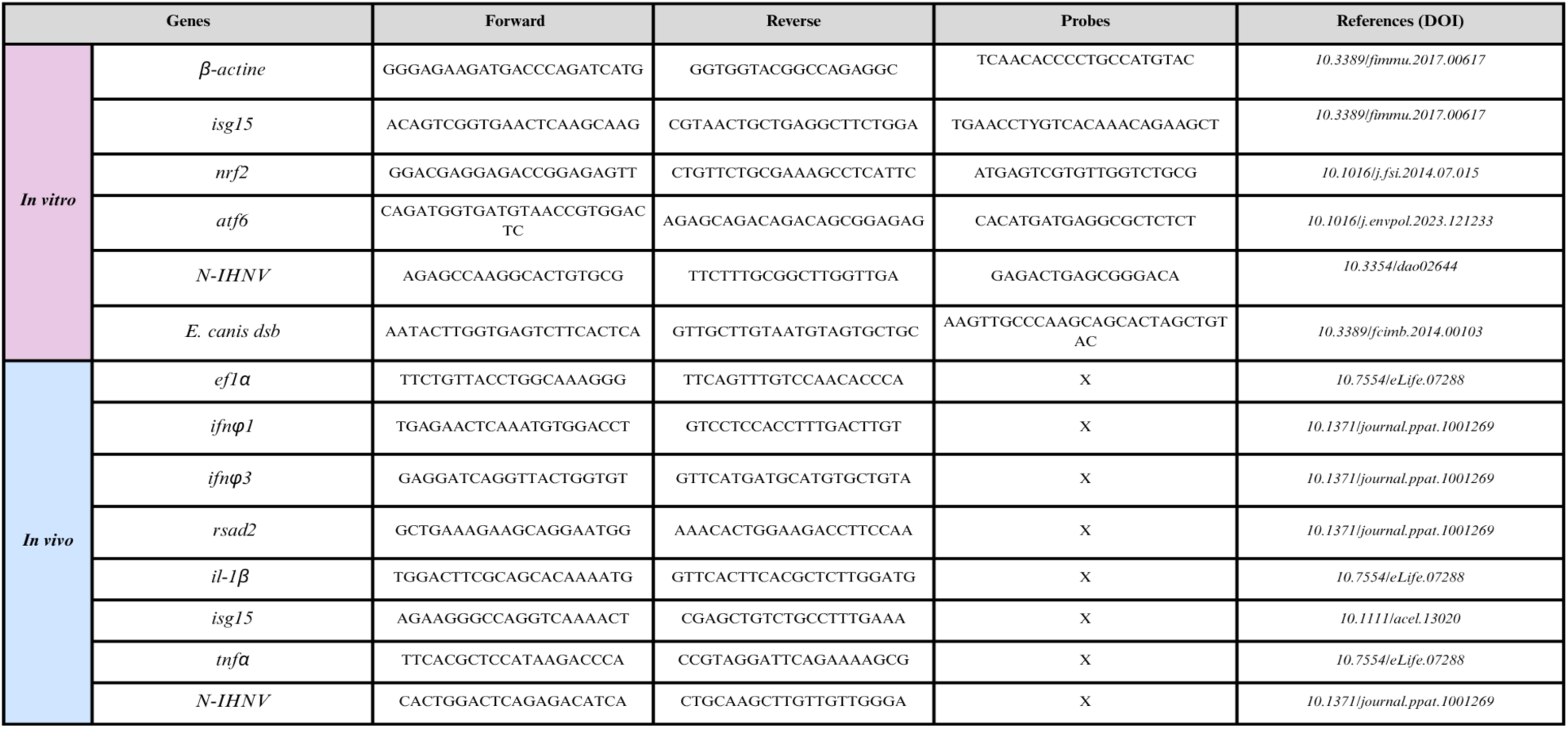
Primer and probe sequences used for RT-qPCR analyses in *in vitro* and *in vivo* experiments. The table lists the forward and reverse primer sequences, TaqMan probe sequences (when applicable), and the corresponding literature references. *β-actin* (*in vitro*) and *ef1α* (*in vivo*) were used as reference genes for normalization.

#### 2.6.1. From cell culture

Total RNA was extracted from 100 µL of viral inoculum, cell supernatants and cell lysates at different time points (0, 24, 48 and 72 hpi). Viral inoculum and cell supernatants were processed using the NucleoSpin RNA Virus Kit (Macherey-Nagel, France) according to the manufacturer’s instructions, cell lysates were homogenized and applied to gDNA Eliminator spin columns to remove genomic DNA before extraction using the RNeasy Plus Mini Kit (Qiagen, France). Three biological replicates were extracted for each strain, kinetic point and temperature studied. To monitor the efficiency of the analytical workflow, a known amount of the exogenous RNA bacteriophage MS2 (Ninove et al., 2011) was added into the samples prior to lysis. All oligonucleotides were validated by RT-qPCR prior to use on microfluidic high-throughput qPCR (Standard BioTools, USA). TaqMan probes were labeled with 6-FAM and BHQ1 (Black Hole Quencher). Table 2 lists the primers/probes used to quantify the expression of *n_IHNV_,* the viral nucleoprotein gene in viral inoculum and cell supernatant, *isg15*, an interferon-stimulated gene and a ubiquitin-like protein traditionally known for its role in ISGylation; *nrf2*, a transcription activator that binds to antioxidant response (ARE) elements; and *atf6*, an endoplasmic-reticulum (ER)-membrane-bound transcription factor activated by ER stress-induced proteolysis, in cell lysates.

Total RNA was reverse-transcribed into cDNA using the RevertAid H Minus First Strand cDNA Synthesis Kit (Thermo Fisher Scientific, USA) according to the manufacturer’s instructions. Briefly, 9–11 µL of treated and diluted RNA was reverse-transcribed using random hexamer primers. Reverse transcription was performed at 25°C for 5 min, 42°C for 60 min and 70°C for 5 min. The resulting cDNA was stored at −20°C until further analysis. Prior to high-throughput qPCR, cDNA samples were pre-amplified using the TaqMan PreAmp Master Mix (Standard BioTools, USA) according to the manufacturer’s instructions. Primers were pooled at a final concentration of 200 nM each. A recombinant pBluescript II SK+ plasmid containing the *dsb* gene from *Ehrlichia canis* was added to the pre-amplification mix as an inhibition control (IC), as previously described by Michelet et al. (2014). Each 5 µL pre-amplification reaction contained 1 µL TaqMan PreAmp Master Mix, 1.25 µL pooled primers, 0.25 µL nuclease-free water, 1 µL of the *dsb,* inhibition control (25 copies/µL), and 1.5 µL cDNA. Pre-amplification consisted of an initial denaturation step at 95°C for 10 min, followed by 14 cycles of 95°C for 15 s and 60°C for 4 min. Pre-amplified products were subsequently diluted 1:5 and stored at −20°C until analysis.

High-throughput real-time PCR was performed using two 192.24 Dynamic Array IFCs on a BioMark™ HD real-time PCR system (Standard BioTools, USA). TaqMan assays were performed using PerfeCTa qPCR ToughMix, Low ROX (QuantaBio, USA). Each sample mix contained 2 µL of PCR master mix, 0.2 µL of 20× GE Sample Loading Reagent (Standard BioTools, USA), and 1.8 µL of diluted pre-amplified cDNA. Each assay mix contained 2 µL of 2× Assay Loading Reagent and 2 µL of 20× oligonucleotide mix, corresponding to 9 µM of each primer and 2 µM of probe. Three µL of each sample mix and assay mix were loaded into the respective inlets of the Dynamic Array IFC. Amplification consisted of 2 min at 50°C, 10 min at 95°C, followed by 40 cycles of 95°C for 15 s and 60°C for 1 min. Each array included no-template controls (H₂O), pre-amplification negative controls (H₂O pre-amp), positive controls for each target, and the *dsb*-based inhibition control. Following pre-amplification, the approximate limit of detection was Ct 24–25. Relative gene expression was calculated using the Pfaffl method (Michael W. Pfaffl, 2001), with *β-actin* serving as the reference gene.

#### 2.6.2. From Zebrafish

IHNV-infected and mock-infected larvae were collected in RLT buffer at 0 and 48 hpi, using 3 pools of 5 larvae per condition. RNA was extracted using the RNeasy Plus Mini Kit (Qiagen) following the manufacturer’s protocol. Total RNA was quantified using a NanoDrop 2000c spectrophotometer (Thermo Scientific) and stored at -80°C until use. RT-qPCR was carried out as described by Souto et al (2024). The primers used for RT-qPCR analyses of IHNV load, interferon signaling pathway (*ifn φ1, ifn φ3, rsad2, isg15*) and pro-inflammatory responses (*il1β and tnfα*) are listed in Table 2. Relative gene expression was calculated using the Pfaffl method (Michael W. Pfaffl, 2001), with *ef1a* (eukaryotic translation elongation factor 1 alpha 1 like 2) serving as the reference gene.

### 2.7. Imaging of Zf larvae and image processing

Stereomicroscopic imaging of anesthetized larvae was carried out using a Leica MZ10F stereomicroscope equipped with an external light source (Leica EL6000), a PLAN APO 1× objective, and an MC170HD camera (Leica Microsystems). Images were processed with Fiji (https://imagej.net/software/fiji/). Brightness and contrast of images were adjusted using the same parameters for each condition. For quantification of neutrophil recruitment, ROI (region of interest) were delimited using ROI manager. Mean gray value of the GFP signal inside the ROI was measured per individual. Data were normalized to percentages for plotting. Fifteen larvae were analyzed per condition.

### 2.8. Statistical analysis

Viral titers obtained from cell culture experiments were analyzed using two-way analysis of variance (two-way ANOVA), followed by Tukey’s multiple comparisons test. Statistical analyses of qPCR data and mpx recruitment data were performed using GraphPad Prism software (version 9). Comparisons between two groups were conducted using the non-parametric Mann–Whitney test. Data are presented as mean ± SEM. A *p* value < 0.05 was considered statistically significant, with statistical significance indicated as follows: *p* < 0.05, \**p* < 0.01, \*\**p* < 0.001, and \*\*\**p* < 0.0001.

## 3. Results

### 3.1. Phylogenetic relationship between selected IHNV field isolates

To investigate the potential role of climate warming in shaping the emergence and selection of circulating IHNV variants and explore the relationship between phylogenetic diversity and thermotolerance, six filed isolates were selected originated from the large collection maintained in the WOAH reference laboratory for IHNV and the French National Reference Laboratory (NRL) for regulated fish diseases. The selection was based on several criteria, including genogroup and geographic diversity, the variety of host fish species and their experimentally-determined *in vivo* virulence levels in rainbow trout. The full nucleotide sequences of the six pre-selected IHNV isolates were enriched with some reference strains available on GenBank to create a ML phylogenetic reconstruction including all described genogroups (Figure 1). The N61, 1075b, and SA-22 isolates fell within genogroup E, with a molecular evolution reflecting their chronological isolation dates. Isolates FR0031, RtNag0a and WRAC were positioned into genogroups L, J and M, respectively.

**Figure 1.**
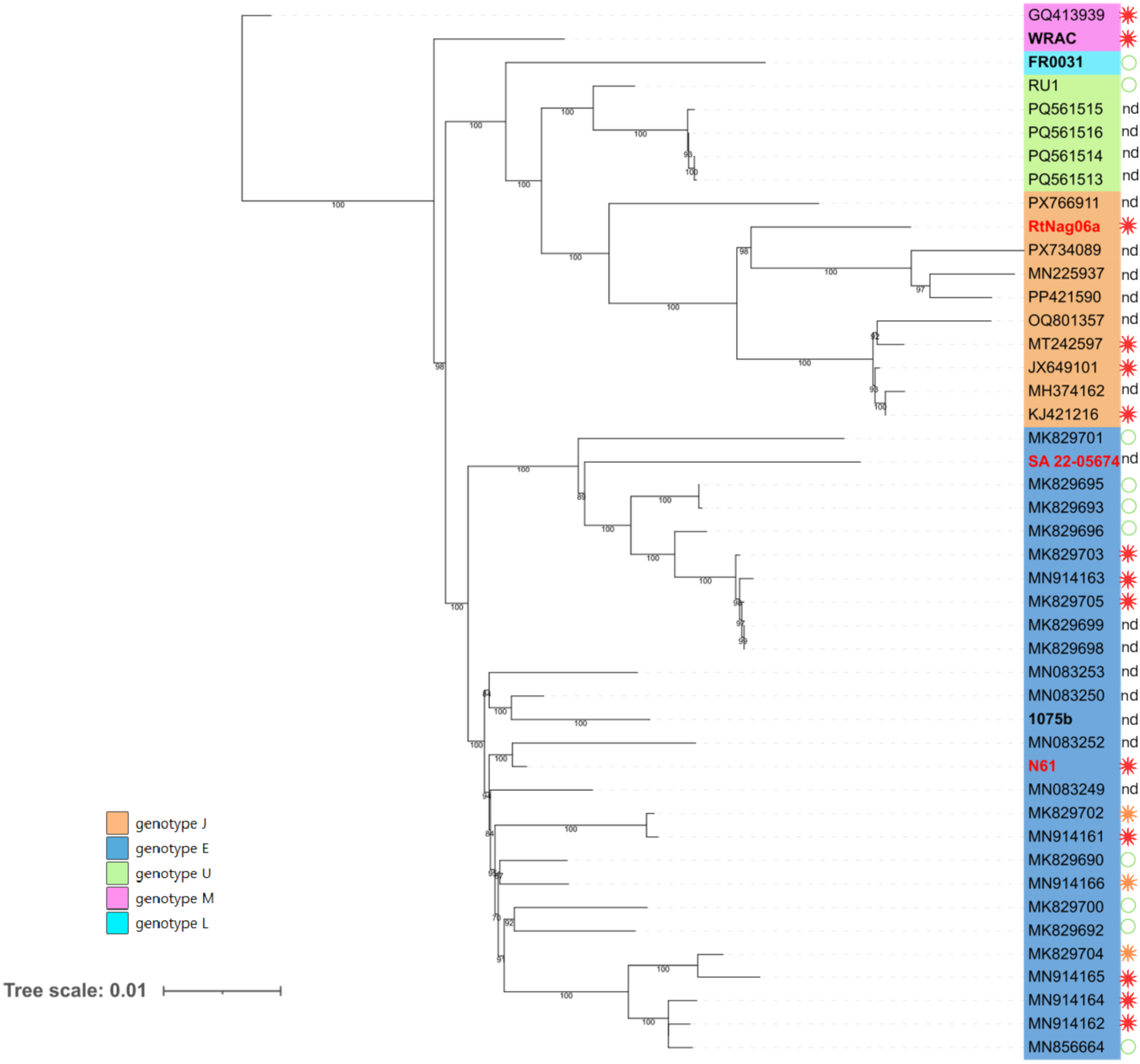
**Maximum likelihood (ML) phylogenetic tree inferred from the complete genome sequences of 47 IHNV isolates**, including the six isolates selected in this study together with representative reference strains retrieved from GenBank and the published literature. The tree was midpoint-rooted and visualized using iTOL (Interactive Tree Of Life). Branch lengths are proportional to the number of nucleotide substitutions per site (scale bar = 0.01 substitutions per site).The six isolates used in this study appear in bold, and the three that underwent more detailed characterization appear in red. Reported IHNV virulence levels in rainbow trout (*Oncorhynchus mykiss*) were classified according to cumulative percent mortality (CPM) as follows: green circles indicate low virulence (CPM ≤ 40%), orange asterisks indicate moderate virulence (40% < CPM < 70%), and red asterisks indicate high virulence (CPM ≥ 70%) (Abbadi et al., 2021). Isolates for which virulence data were not available are indicated as nd (not determined).

### 3.2. Assessment of the thermotolerant profile of selected IHNV field isolates

Replication of these isolates was monitored at 14°C (optimal temperature of growth), 20, 22, and 24 °C (considered as suboptimal temperatures) throughout the 8 days following infection. Pre-screening assays were carried out on EPC cells to distinguish their thermotolerance profiles. All isolates showed similar capacities of replication on EPC cells at 14°C, with a progressive increase in the initial amount of infectious virus (inoculum) starting at 2 days post infection (dpi) and reaching 3.0 to 3.5 Log_10_ TCID_50_*/*mL increase at 8 dpi (Table 3). At 20 °C, the kinetics were accelerated for isolate SA-22, with earlier and more intense replication (increases of +1.6 and +2.9 Log_10_ TCID_50_*/*mL at 1 and 2 dpi, respectively, compared to 14°C). This isolate showed at least a level of replication 1.7 Log_10_ more important than the other strains at 8 dpi. Isolate RtNag0a also had an earlier replication (+1.2 Log_10_ TCID_50_*/*mL at 2 dpi) but without exceeding the maximum level observed at 14°C. Isolate 1075b displayed comparable replication kinetics at both 14 and 20°C between 2 and 6 dpi. Isolates N61 and WRAC showed weaker replication at 20°C, and this situation was more marked for strains FR0031. At 22°C, a significant replication was observed for SA-22 and, to a much lesser extent, for RtNag0a. No isolate replicated at 24°C. These results allowed for the selection of 3 IHNV isolates for their thermotolerance profiles to perform in-depth analyses: SA-22 and RtNag0a which appear to have a higher replication potential than others at suboptimal temperatures of 20 and 22°C, and N61 which is the reference strain used by the French NRL.

**Table 3.** Synthesis of the replication capacities of the six IHNV isolates as a function of temperature. The values indicated (in Log₁₀ TCID₅₀/mL) correspond to the difference between the viral titer measured in the mixture of cell lysate and culture supernatant at the end of the experiment and the initial inoculum titer (3.7 Log₁₀ TCID₅₀/mL) used to infect the cells. The background color represents the level of viral replication under each condition, with colors progressively shifting toward red as replication increases. Values on a white background indicate no detectable replication. The letter preceding each isolate name corresponds to its genotype. Measurements were performed in triplicate for isolates SA-22, RtNag0a, and N61, with standard errors of measurement (SEM) ≤ 0.37, 0.70, and 0.51, respectively, and once for the other isolates.

|  | **14°C** |  |  |  |  | **20°C** |  |  |  |  | **22°C** |  |  |  |  | **24°C** |  |  |  |  |
|---|---|---|---|---|---|---|---|---|---|---|---|---|---|---|---|---|---|---|---|---|
| *Days post-infec* | 1 | 2 | 3 | 6 | 8 | 1 | 2 | 3 | 6 | 8 | 1 | 2 | 3 | 6 | 8 | 1 | 2 | 3 | 6 | 8 |
| **E-SA-22** | 0,0 | 1.3 | 2.7 | 3.7 | 3.3 | 1.6 | 4.2 | 3.8 | 3.6 | 4.3 | 2.2 | 2.5 | 3.0 | 1.5 | 1.7 | 0.0 | 0.0 | 0.0 | 0.0 | 0.0 |
| **E-N61** | 0,1 | 1.9 | 2.9 | 3.3 | 3.4 | 0.2 | 2.2 | 2.2 | 1.8 | 2.1 | 0.2 | 0.0 | 0.0 | 0.0 | 0.0 | 0.0 | 0.0 | 0.0 | 0.0 | 0.0 |
| **E-1075b** | 0,0 | 0.5 | 1.5 | 3.0 | 3.5 | 0.0 | 1.0 | 1.7 | 3.2 | 0.9 | 0.0 | 0.0 | 0.0 | 0.0 | 0.0 | 0.0 | 0.0 | 0.0 | 0.0 | 0.0 |
| **J-RtNag0a** | 0,0 | 0.9 | 2.1 | 2.9 | 3.0 | 0.0 | 2.1 | 2.5 | 2.2 | 2.6 | 0.0 | 0.0 | 1.0 | 1.0 | 0.7 | 0.0 | 0.0 | 0.0 | 0.0 | 0.0 |
| **L-FR0031** | 0,0 | 0.7 | 1.5 | 3.5 | 3.5 | 0.2 | 0.2 | 0.0 | 1.4 | 0.4 | 0.0 | 0.0 | 0.0 | 0.0 | 0.0 | 0.0 | 0.0 | 0.0 | 0.0 | 0.0 |
| **M-WRAC** | 0,0 | 1.5 | 3.2 | 4.0 | 3.5 | 0.5 | 1.0 | 2.2 | 2.7 | 2.4 | 0.0 | 0.0 | 0.0 | 0.0 | 0.0 | 0.0 | 0.0 | 0.0 | 0.0 | 0.0 |

### 3.3. In vitro characterization of viral isolates exhibiting the highest replicative thermoprofile

#### 3.3.1. Infectious particle release kinetics and phenotypes

We next characterized in details the replicative thermoprofiles of the three selected isolates N61, SA-22 and RtNag0a by analyzing the viral productions in the supernatant of EPC over a 72 hours period post infection at three temperatures: 14, 20 and 22°C. At 14°C, viral titers increased progressively in the supernatant for all three isolates (Figure 2A). SA-22 exhibited the highest viral production, reaching 6.6 Log_10_ TCID_50_*/*mL at 72 hpi. N61 and RtNag0a showed slower replication kinetics. Although RtNag0a displayed slightly higher titers than N61 up to 48 hpi, both isolates reached comparable levels by 72 hpi, with final titers of approximately 5.2 and 5.5 Log_10_ TCID_50_*/*mL, respectively. At 20°C, replication kinetics differed more markedly between isolates. SA-22 showed a rapid increase in viral titer, reaching approximately 7.5 Log_10_ TCID_50_*/*mL by 48 hpi and remaining stable thereafter. N61 and RtNag0a also displayed increasing titers over time but reached lower final values of approximately 5.4 and 5.8 Log_10_ TCID_50_*/*mL, respectively. At 22°C, SA-22 again showed the highest replication efficiency, with viral titers increasing rapidly to approximately 6.8 Log_10_ TCID_50_*/*mL by 48 hpi. RtNag0a exhibited intermediate replication kinetics, reaching approximately 4.9 Log_10_ TCID_50_*/*mL at 72 hpi, whereas N61 showed the lowest viral production, with titers remaining below 3.7 Log_10_ TCID_50_*/*mL throughout the experiment. Overall, viral titers increased over time for all isolates, although the magnitude of replication depended on both the viral isolate and incubation temperature. SA-22 consistently reached the highest viral titers at all temperatures tested, while the replication of N61 was more strongly reduced at 22°C. RtNag0a displayed an intermediate replication phenotype across the three temperatures tested.

**Figure 2.**
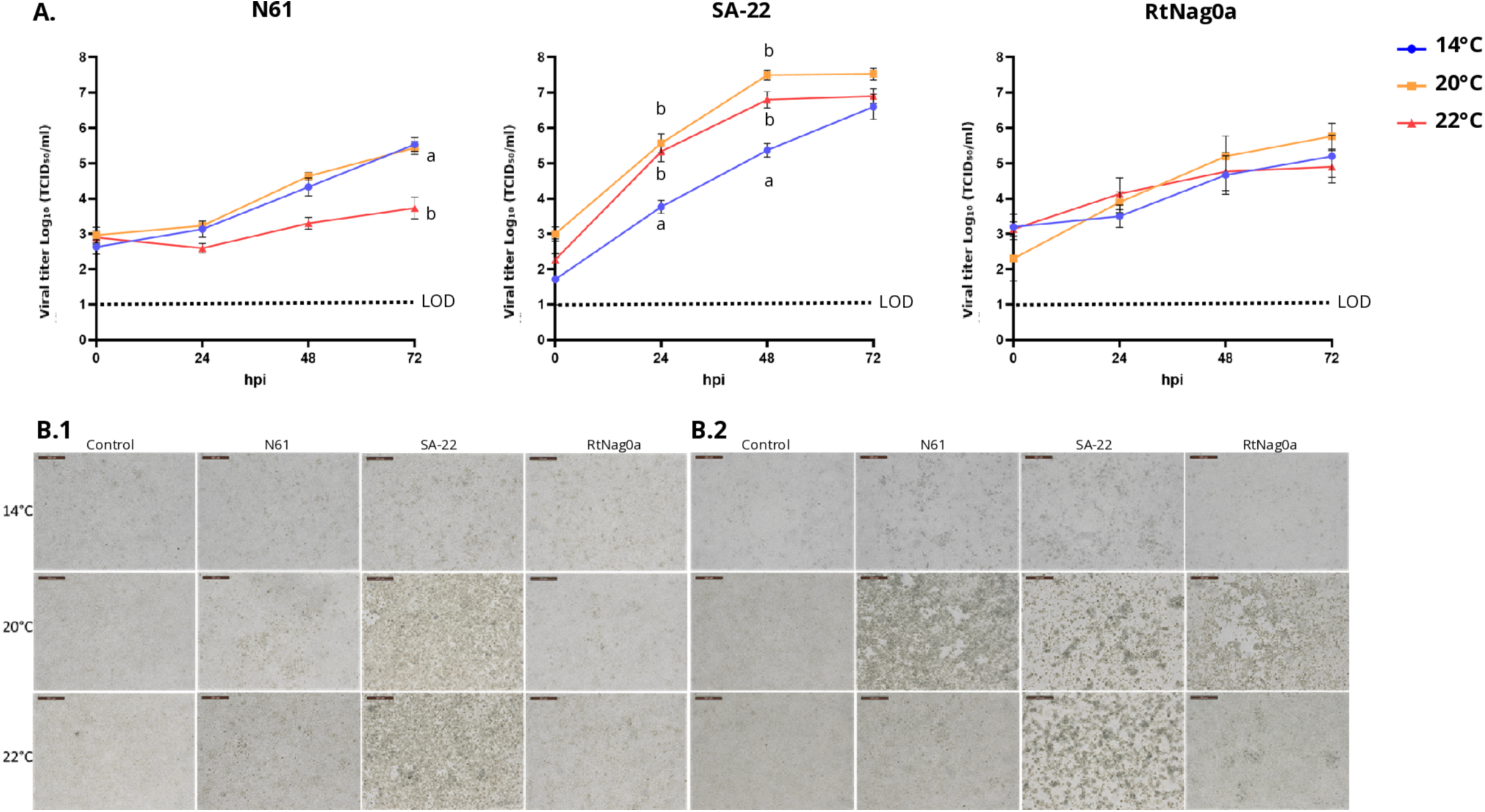
Temperature dependent of IHNV replication on EPC. **A.** Number of infectious particles (in Log_10_ TCID_50_/mL) were measured from culture supernatants collected at the indicated time points following infection with N61, SA-22 and RtNag0a at an MOI of 0.1 at 14, 20 and 22°C. Viral titers were determined by plaque assay (5 measurements per point for each strain). Average values are shown with the standard error of measurement (SEM). The dotted line indicates the limit of detection (LOD). The different letters indicate statistically significant differences between viral titers with regards to temperature (P<0.05). **B.** Representative microscopy images of infected cell monolayers at 48 (**B.1**) and 72 (**B.2**) hpi at 14, 20 and 22°C at ×200 total magnification (20× objective and 10× ocular lens). Scale bar = 200 µm.

We then assessed virus-induced cytopathic effects (CPE) comparing 14, 20 and 22°C from 48 to 72 hpi. At 14°C, all isolates produced only mild CPE, with minimal progression between 48 and 72 hpi compared with uninfected cell monolayers (Figure 2B). At 20°C, severe CPE was already evident at 48 hpi in SA-22 infected cells, while N61 induced moderate CPE and RtNag0a remained mild. By 72 hpi, all isolates produced severe CPE at this temperature. Increasing the temperature to 22 °C did not alter the phenotypes observed for the 20°C condition at 48 hpi, but by 72 hpi, only SA-22 induced a complete CPE, whereas N61 and RtNag0a maintained a stable mild effect. Overall, these observations show that SA-22 induced the strongest cytopathic response particularly at 20–22 °C, whereas N61 and RtNag0a only triggered mild CPE at 14 and 22°C but reached a severe phenotype at 20°C. Thus, the replication capacity of these isolates is not directly correlated with the extent of CPE during the observation period (until 72 hpi) as all isolates replicated efficiently at 14 °C without inducing major alteration of the cell monolayers.

#### 3.3.2. Evolution of the number of intracellular and released genome copies

To further characterize viral replication dynamics at different temperatures, viral RNA levels were monitored over time in intracellular and released fractions.

In cell culture supernatants, viral RNA progressively increased over time for all three isolates, although the magnitude of replication differed between isolates and temperatures (Figure 3A). N61 showed a relatively limited increase, with the highest viral RNA level observed at 20°C (7.63 × 10¹⁰ RNA copies/mL at 72 hpi), while replication was strongly reduced at 22°C (6.9 × 10⁹ RNA copies/mL). In contrast, SA-22 showed a marked increase in viral RNA, with 9.26 × 10¹¹, 4.14 × 10¹² and 2.34 × 10¹² RNA copies/mL at 14, 20 and 22°C respectively, at 72 hpi. RtNag0a also showed substantial viral RNA production, particularly at 20°C, reaching 1.27 × 10¹² RNA copies/mL at 48–72 hpi, compared to 3.45 × 10¹¹ and 2.36 × 10¹¹ RNA copies/mL at 14 and 22°C, respectively.

**Figure 3.**
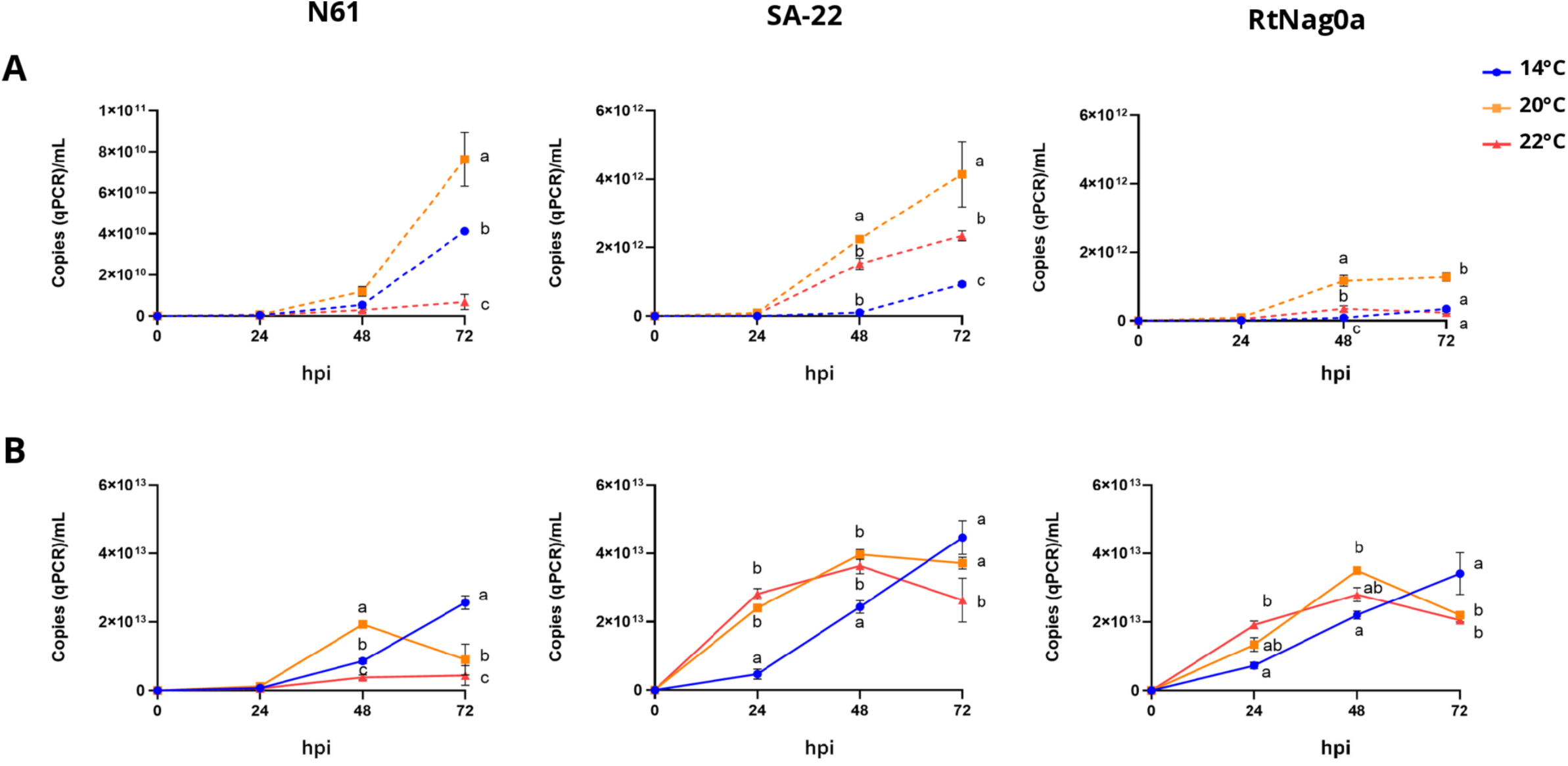
Intra and extracellular viral RNA load following EPC infection with the three selected IHNV isolates. Viral RNA levels were quantified by RT-qPCR for N61, SA-22 and RtNag0a at 14°C, 20°C and 22°C. **A.** Viral copy numbers measured in culture supernatants at 0, 24, 48 and 72 hpi. **B.** Viral copy numbers measured in cell lysates at the same time points. Data are presented as mean ± SEM (n = 3 biological replicates per strain, time point and temperature). Statistical analyses were performed using two-way ANOVA followed by Tukey’s multiple-comparisons test for panels A and B.

A clear accumulation of viral RNA was also observed in cell lysates throughout infection (Figure 3B). N61 reached its highest intracellular viral RNA level at 14°C (2.57 × 10¹³ RNA copies/mL at 72 hpi), whereas lower levels were detected at 22°C (4.39 × 10¹² RNA copies/mL) and 20°C (9.00 × 10¹² RNA copies/mL). SA-22 and RtNag0a showed faster kinetics, both reaching high intracellular RNA levels at all temperatures, with maximum values of approximately 4.46 × 10¹³ and 3.5 × 10¹³ RNA copies/mL, respectively. Overall, these data demonstrate productive replication of the three IHNV isolates across the tested temperature range, with distinct temperature-dependent replication profiles and generally higher viral RNA accumulation for SA-22 and RtNag0a than for N61. It is interesting to note that the intracellular/extracellular ratios of viral RNA are lower for the SA-22 and RtNag0a isolates compared to the N61 strain.

#### 3.3.3. Effect of temperature and viral isolate on cell responses

The influence of incubation temperature on EPC cellular machinery and the response to IHNV infection was investigated by measuring the expression profiles of *isg15*, *nrf2*, and *atf6*. These genes were selected to investigate three complementary cellular pathways, interferon-mediated antiviral response *(isg15)*, response to oxidative stress *(nrf2)*, and unfolded protein response associated with endoplasmic reticulum *(atf6)*.

Expression of *isg15*, *nrf2*, and *atf6* was generally induced upon infection with the three thermotolerant IHNV isolates, with responses varying according to both time post-infection and temperature (Figure 4). At 14°C, *isg15* expression showed a significant increase at 72 hpi (Figure 4A). At 20°C, expression significantly increased over the course of the infection, in a more nuanced way for the SA-22 isolate. At 22°C, *isg15* expression significantly increased after 48 hpi. These results indicate differences in the temporal profile of *isg15* induction across temperature conditions.

**Figure 4.**
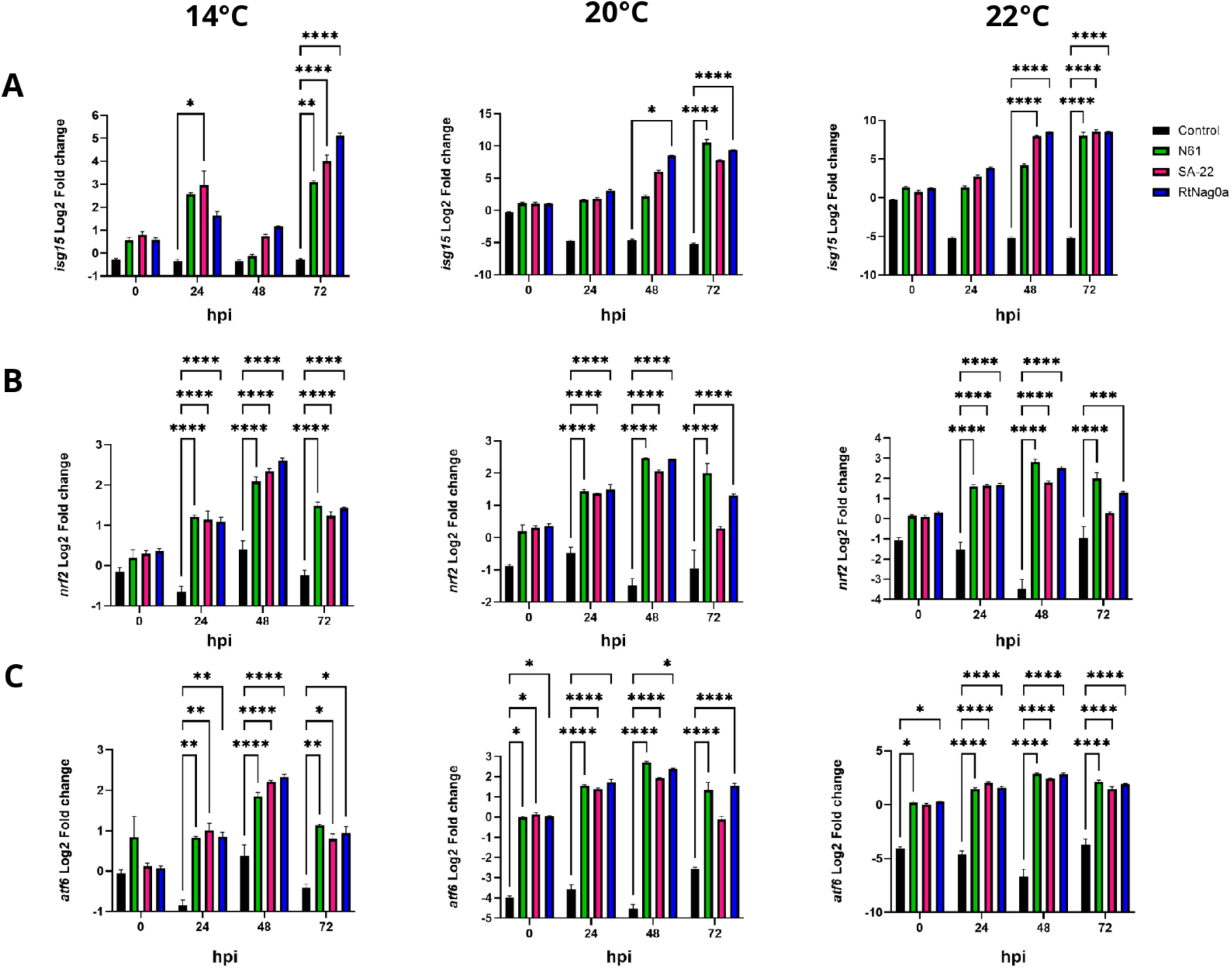
Gene expression profiles following infection with the three selected IHNV isolates. Gene expression was quantified by high-throughput real-time PCR in response to infection with N61, SA-22, RtNag0a and mock at the indicated temperatures and time points (0, 24, 48 and 72 hpi). **(A)** *isg15*, **(B)** *nrf2* and **(C)** *atf6* expression levels. Data are expressed as log₂ fold change relative to the control condition and presented as mean ± SEM. Statistical analyses were performed using two-way ANOVA followed by Tukey’s multiple-comparisons test. Significant differences are indicated by asterisks (*p < 0.05, **p < 0.01, ***p < 0.001, ****p < 0.0001).

*nrf2* expression (Figure 4B) increased significantly over the course of infection under all three temperature conditions, although the temporal pattern varied depending on the temperature and viral isolate. At 14°C, *nrf2* expression displayed a significant time-dependent increase, except between 24 and 72 hpi where levels remained stable. At 20°C, N61 showed a significant temporal rise. For SA-22, significant differences were detected between most time points, except between 0 and 72 hpi, where levels remained comparable. For RtNag0a, all pairwise comparisons were significant except the 24–72 hpi interval, which showed no detectable change. At 22°C, N61 and RtNag0a showed significant increases over time, except between 24 and 72 hpi, whereas SA-22 exhibited significant increases between 0 and 24 hpi and between 24 and 48 hpi.

Finally, *atf6* expression (Figure 4C) was induced following infection, with distinct temporal profiles depending on the temperature. At 14°C, *atf6* expression significantly increased from 0 to 48 hpi in all three isolates, followed by a significant decrease at 72 hpi. At 20°C, expression significantly increased progressively up to 48 hpi, before significantly decreasing at 72 hpi to levels comparable to those observed at 0 and 24 hpi. At 22°C, *atf6* expression significantly increased at each time point throughout the infection. Overall, these results indicate a dynamic regulation of *atf6* expression throughout infection, with temporal profiles varying according to temperature and viral isolate.

### 3.4. Replicative thermoprofile of selected field isolates in Rainbow Trout derived cell line

To rule out a potential interspecies barrier affecting the *in vitro* infection assays, we next characterized the replication capacity of N61, RtNag0a and SA-22 isolates in rainbow trout–derived RTG-2 cell line and measured replication kinetics at 14 and 20°C (Table 4), 20°C corresponding to the highest temperature routinely used at the French Reference Laboratory (NRL). As described for EPC cells (Table 3), viral replication was expressed relative to the initial titer of the inoculum over 8 days post infection period. All viral isolates efficiently replicated in RTG-2, with relatively similar viral amplification at 8 dpi, from 3.6 Log_10_ TCID_50_*/*mL for RtNag0a and SA-22 to 4.1 for N61 at 14°C. Notably, N61 showed a gradual increase in amplification from 1 to 8 dpi, whereas replication of RtNag0a and SA-22 was not detectable before 6 dpi. At 14°C, an increase of more than 3 Log10 TCID50/mL relative to the initial inoculum was observed from 2 dpi for N61, compared with 6 dpi for RtNag0a and 8 dpi for SA-22. By 8 dpi at 20°C, N61 and SA-22 achieved comparable viral titers (4.3 and 4.5 log_10_ TCID_50_/mL, respectively), while RtNag0a showed slightly reduced replication efficiency, reaching 3.6 log_10_ TCID_50_/mL. Overall, viral replication kinetics were accelerated and reached higher levels at 20°C compared with 14°C (for N61 and SA-22), contrasting with the results obtained in EPC cells.

**Table 4.**
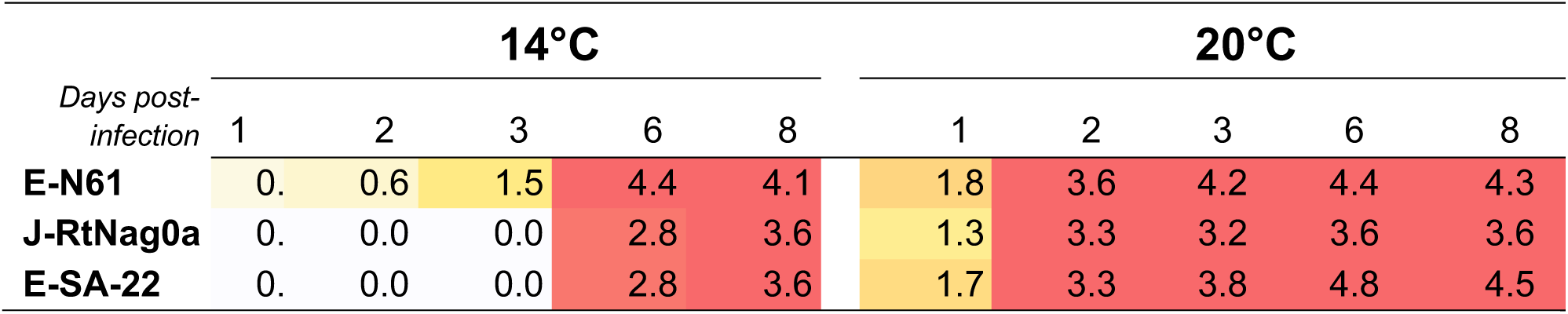
Synthesis of the replication capacities of IHNV isolates N61, SA-22, and RtNag0a on the RTG-2 cell line at 14°C and 20°C. The number of infectious particles was measured in culture supernatants and cell lysates collected at the indicated time points following infection at an MOI of 0.1. The titers indicated (in Log₁₀ TCID₅₀/mL) correspond to the difference between the viral titer measured at each time point and the initial inoculum titer (3.7 Log₁₀ TCID₅₀/mL) used to infect the cells. Measurements were performed in triplicate, with standard errors (SEM) ≤ 0.44, 0.79, and 0.63 for isolates N61, SA-22, and RtNag0a, respectively. The background color represents the level of replication under each condition, with colors progressively shifting toward red as replication increases. Values on a white background indicate no detectable replication.

|  | **14°C** |  |  |  |  |  | **20°C** |  |  |  |  |
|---|---|---|---|---|---|---|---|---|---|---|---|
| *Days post-infection* | 1 | 2 | 3 | 6 | 8 |  | 1 | 2 | 3 | 6 | 8 |
| **E-N61** | 0. | 0.6 | 1.5 | 4.4 | 4.1 |  | 1.8 | 3.6 | 4.2 | 4.4 | 4.3 |
| **J-RtNag0a** | 0. | 0.0 | 0.0 | 2.8 | 3.6 |  | 1.3 | 3.3 | 3.2 | 3.6 | 3.6 |
| **E-SA-22** | 0. | 0.0 | 0.0 | 2.8 | 3.6 |  | 1.7 | 3.3 | 3.8 | 4.8 | 4.5 |

### 3.5 Genetic relationships and putative determinants of thermotolerance in IHNV

To further characterize the genetic relationships among IHNV field isolates harboring thermotolerance phenotypes on EPC, we conducted a phylogenetic analysis based on full-length IHNV G-gene sequences. The six isolates of this study were compared with genotype E strains from Italy and Denmark previously classified by Marsella et al. (2025) as thermotolerant (67-1, DK) or thermosensitive (217/A, 224-2, 354). *In vitro* thermo-adapted IHNV25 variant derived from a genotype M strain (Fauvet *et al*. submitted) and italian IHNV/143/14 isolate reported as low virulent in rainbow trout at 10°C were also included in this comparison (Figure 5). Interestingly, isolate SA-22 was positioned within a clade containing isolates IHNV/DK, 143/14, and 67-1 (Figure 5A). This placement groups SA-22 with isolates previously described as thermotolerant, suggesting potential genetic determinants associated with its *in vitro* phenotype.

**Figure 5.**
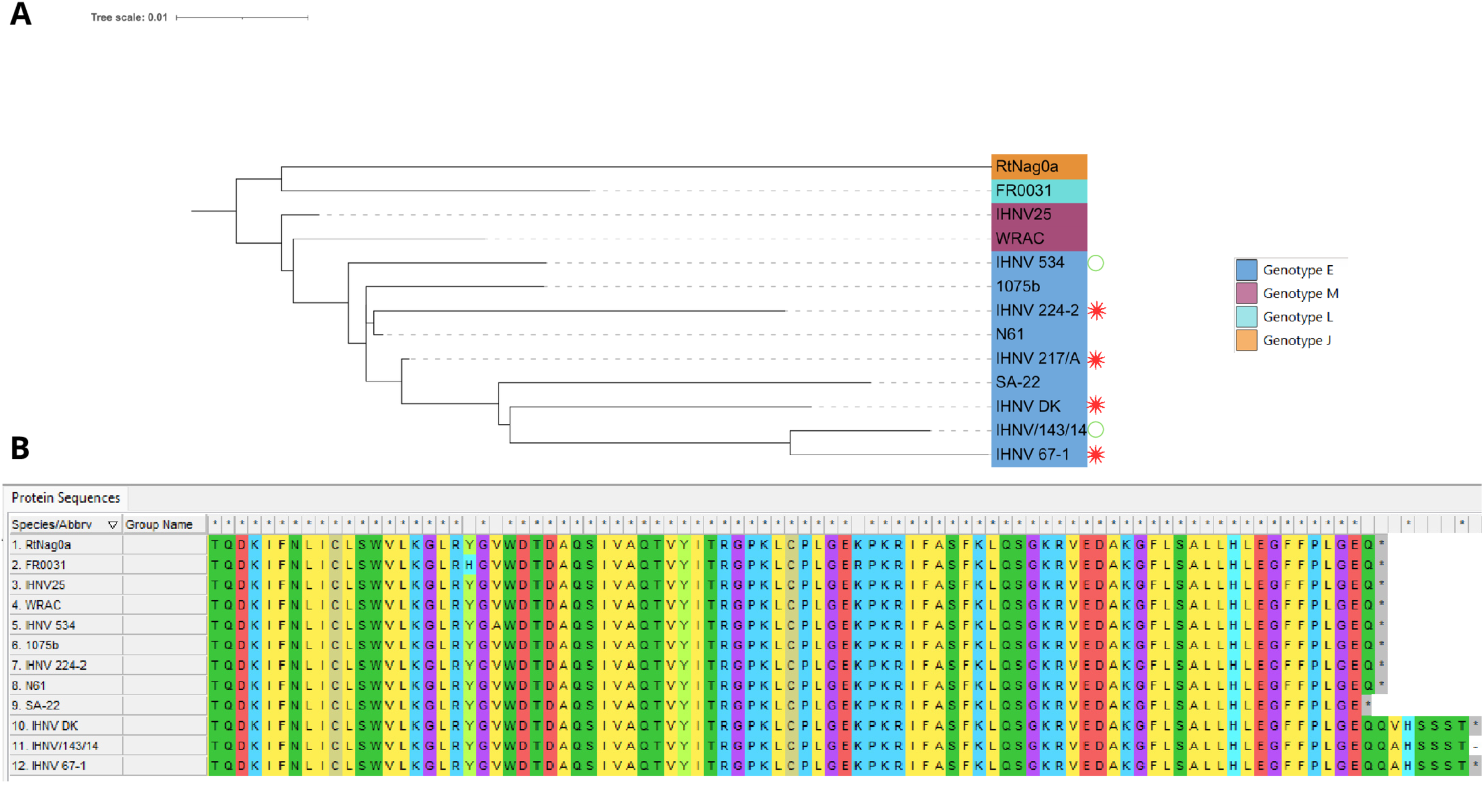
Phylogenetic relationships and comparison of the C-terminal region of the L protein of 12 IHNV isolates. **(A)** Maximum likelihood (ML) phylogenetic tree inferred from the L protein sequences of the 12 IHNV isolates. The tree was midpoint-rooted and visualized using iTOL (Interactive Tree Of Life). Branch lengths are proportional to the number of amino acid substitutions per site (scale bar = 0.01 substitutions per site). Isolates are colored according to their genotype: E, J, M, and L. Reported IHNV virulence levels in rainbow trout (*Oncorhynchus mykiss*) were classified according to cumulative percent mortality (CPM) as follows: green circles indicate low virulence (CPM ≤ 40%), orange asterisks indicate moderate virulence (40% < CPM < 70%), and red asterisks indicate high virulence (CPM ≥ 70%). **(B)** Multiple sequence alignment of the C-terminal region of the L protein from the 12 IHNV isolates. Amino acid sequences are shown using the one-letter amino acid code. Asterisks (*) indicate stop codons.

An extended C-terminal region of the L protein, previously proposed to be associated with thermal adaptation, was observed in IHNV DK, IHNV 143/14, and IHNV 67-1 (Marsella et al., 2025) (Figure 5B). This 7 amino-acid insertion was absent from all other isolates. In contrast, SA-22 harbored a premature stop codon in this terminal region, which truncates the L polymerase by one amino acid, suggesting an isolate-specific structural variation within the C-terminal domain of the L protein.

### 3.6. In vivo investigations of thermotolerant IHNV isolates in Zf larvae

#### 3.6.1. Temperature-dependent virulence of selected IHNV field isolates

To evaluate the virulence of selected field IHNV isolates, Zf larvae were challenged intravenously at 3 dpf, they were maintained at either 21.5°C or 23.5°C and monitored daily for survival throughout the experimental period (Figure 6A). Mortality in mock-infected larvae remained below 10% throughout the experimental period. At 21.5°C, N61 induced an important decrease in survival between 72 and 96 hpi, culminating in 50% mortality at 96 hpi. By comparison, SA-22 and RtNag0a led to earlier mortality events, evident from 48 hpi and exceeding 90% by 96 hpi. A 2° elevation in incubation temperature resulted in a survival rate of N61-infected fish to nearly 80%. In contrast, survival remained unchanged for larvae infected with SA-22 or RtNag0a, with mortality rates remaining at approximately 90% at both temperatures. IHNV infection triggered pathogenicity assessed by clinical scoring (Figure 6B). Clinical signs appeared at 48 hpi regardless of viral isolate or incubation temperature. Pericardial oedema was the predominant clinical sign and was frequently accompanied by exophthalmos and/or hemorrhages. While these lesions occurred at both temperatures in all infected groups, the concomitant occurrence of pericardial oedema and exophthalmos was detected exclusively at 23.5°C (Figure 6B.2) for all three viral isolates, suggesting a temperature-dependent increase in clinical severity.

**Figure 6.**
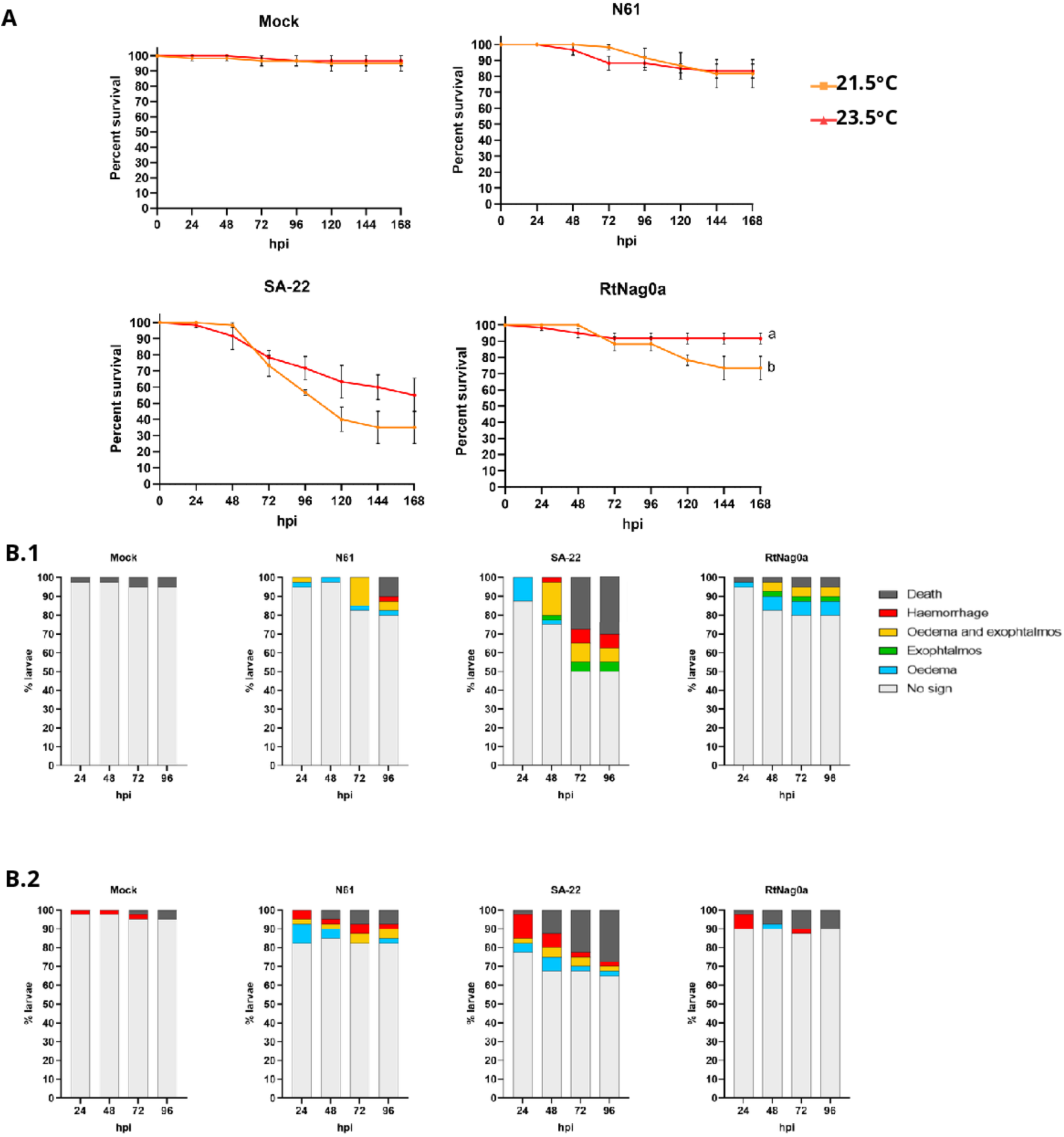
Mortality and clinical signs following intravenous injection of Zf larvae at 21.5°C and 23.5°C with isolates N61, RtNag0a and SA-22. **A.** Cumulative mortality was followed during 168 hpi. Results are expressed as survival rates (%) ± SEM, with assays (n=3) done at independent times. The lowest temperature is represented in orange, the highest in red. **B.** Representative clinical signs observed during the experiment according to the water temperature: 21.5°C (B.1) and 23.5°C (B.2).

#### 3.6.2. Immune responses of Zf larvae induced by IHNV infection

We next examined virus-induced immune responses in whole Zf larvae at 48 hpi using RT-qPCR, focusing on the SA-22 and RtNag0a isolates at 21.5°C (Figure 7). The type I interferon response was assessed by measuring the expression of *ifnφ1*, *ifnφ3*, *rsad2*, and *isg15*. For SA-22, infection resulted in significant upregulation of *ifnφ1*, *ifnφ3*, and *isg15* expression compared with mock-infected larvae. The induction of *ifnφ3* was the most pronounced, while *ifnφ1* expression increased approximately fivefold. Although *rsad2* expression also increased following infection, this difference was not statistically significant (Figure 7A). Pro-inflammatory responses were also induced following SA-22 infection, with significant upregulation of both *il1β* and *tnfα*. The increase in *il1β* expression was more pronounced than that of *tnfα*. In contrast, RtNag0a induced a more limited interferon response, with significant upregulation of *isg15* but no significant changes in *ifnφ1*, *ifnφ3* or *rsad2* expression. RtNag0a infection also resulted in significant upregulation of *il1β* (Figure 7B) without changes of *tnfα*. Finally, viral *N* gene expression significantly increased at 48 hpi for both isolates, confirming viral replication under these experimental conditions.

**Figure 7.**
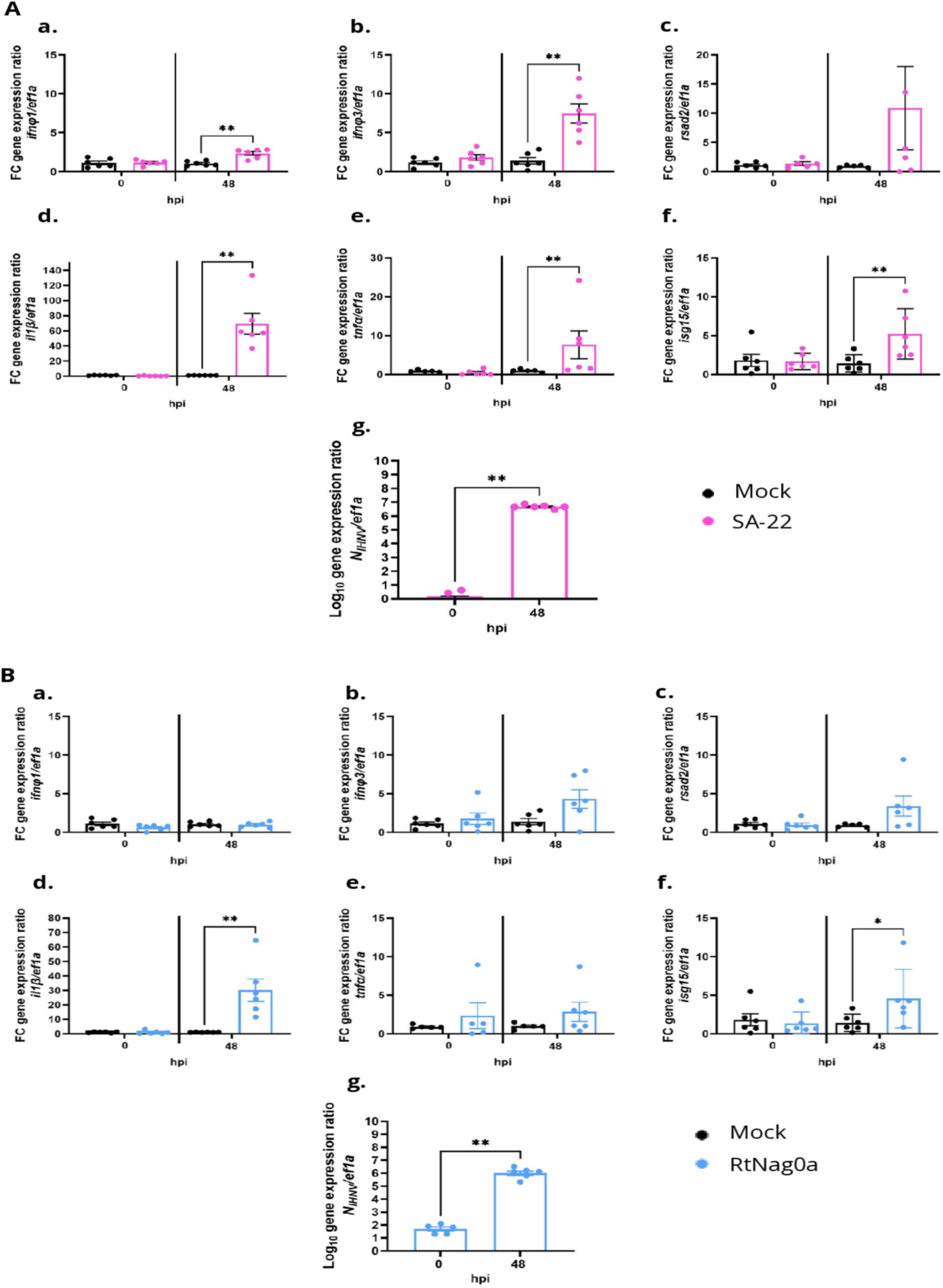
Host and viral gene expression analysis following IV injection of Zf larvae at 21.5°C. Gene expression levels were measured by RT-qPCR at 0 h and 48 hpi at 21.5°C in fish injected with the SA-22 (**A**) and RtNag0a (**B**) isolates, with a mock-injected group as a negative control. (a) *ifn φ1*, (b) *ifn φ3*, (c) *rsad2*, (d) *il1β*, (e) *tnfα*, (f) *isg15*, and (g) viral *n* gene quantification. Panels (a–f) correspond to host immune-related genes, while panel (g) shows viral gene quantification. *ef1α* was used as a reference gene to normalize host gene expression levels. All experiments were performed in duplicate. Statistical significance was assessed using the Mann–Whitney test.

#### 3.6.3. Thermotolerant IHNV field strains successfully infect Zf larvae following waterborne exposure

Based on these results, we subsequently examined whether immersion of Zf larvae in viral suspensions was sufficient to initiate infection, as this exposure method has previously been reported to be ineffective for studying IHNV infections (Ludwig et al., 2011). At 21.5°C, larvae challenged with N61 and RtNag0a progressively died from infection starting at 72 and 96 hpi, respectively, with a cumulative mortality of approximately 20% at 144 hpi (20% for N61 and 27% for RtNag0a) (Figure 8). Exposure to SA-22 used at a concentration 20 times higher culminated in about 70% mortality at 144 hpi. Increasing the incubation temperature to 23.5 °C slightly reduced the mortality rate for all three isolates. At 168 hpi, SA-22 still induced around 50% mortality. N61 and RtNag0a resulted in only 9 and 17% mortality respectively. Daily monitoring showed that immersion led to the same clinical signs as those observed following injection, including hemorrhages, exophthalmos and oedema (Figure 8B). These symptoms were observed at both incubation temperatures and appeared as early as 24 hpi.

**Figure 8.**
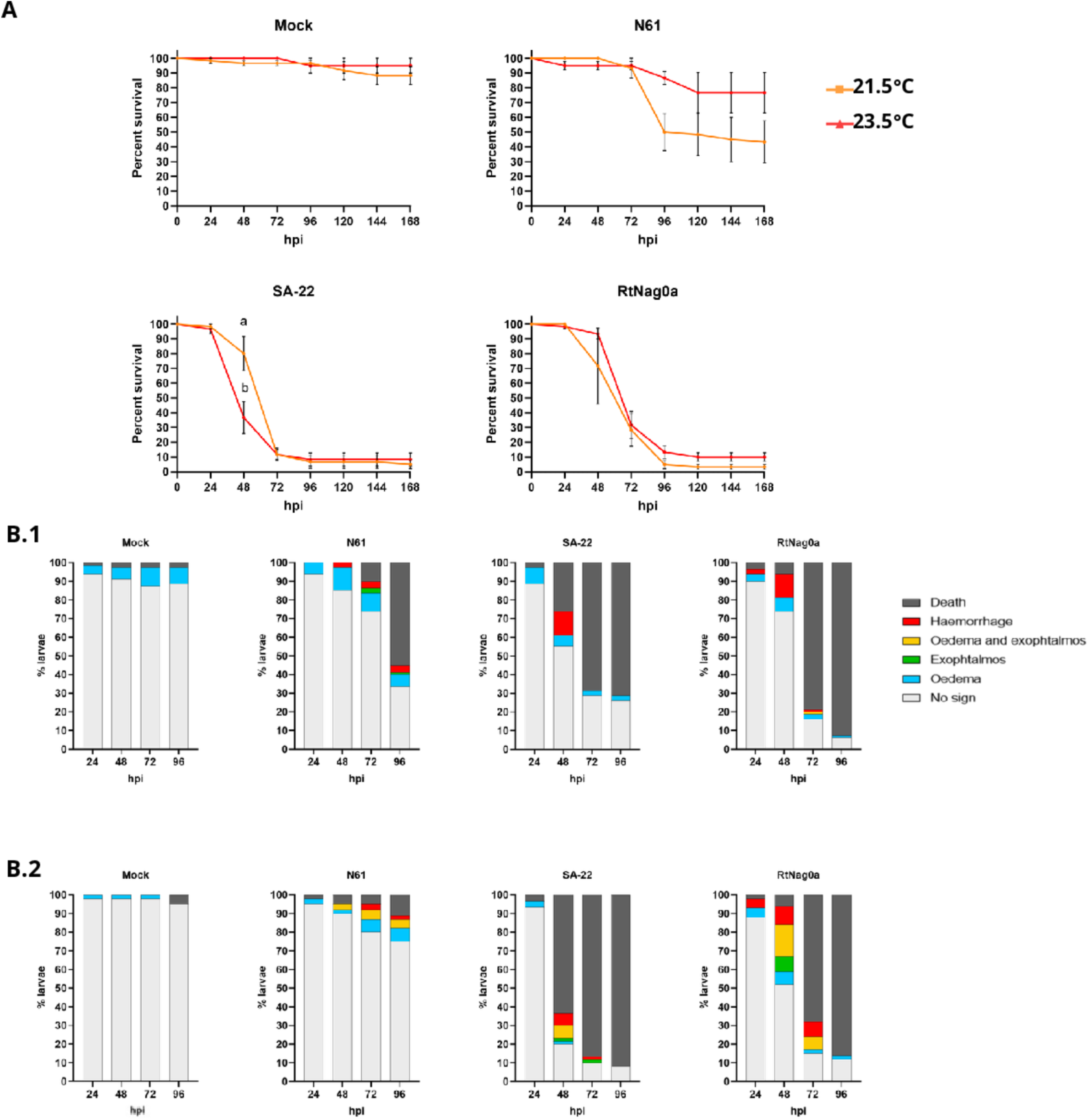
Mortality and clinical signs following infection by immersion of Zf larvae at 21.5°C and 23.5°C with isolates N61, RtNag0a and SA-22. **A.** Cumulative mortality was followed during 168 hpi. Results are expressed as survival rates (%) ± SEM, with assays (n=3) done at independent times. The lowest temperature is represented in orange, the highest in red. **B.** Representative clinical signs observed during the experiment according to the water temperature: 21.5°C (B1) and 23.5°C (B2).

Viral loads were monitored from 6 to 48 hpi following either IV injection or immersion at 21.5 and 23.5°C (Figure 9). In IV injected larvae, no infectious virus was detected during the first 6 hpi at either temperature. Viral production became detectable from 24 hpi and increased markedly by 48 hpi, reaching up to approximately 5.7 Log₁₀ PFU/mL at 21.5°C, compared with lower levels at 23.5°C (Figure 9A). Similar kinetic profiles were observed following immersion, although only one out of four larvae was positive in each time group. This is consistent with the lower mortality rates observed following waterborne challenge (Figure 8A). In contrast, intravenous injection yielded productive infection in all examined larvae producing higher level of infectious viral particles than immersion at 24 and 48 hpi at both temperatures.

**Figure 9.**
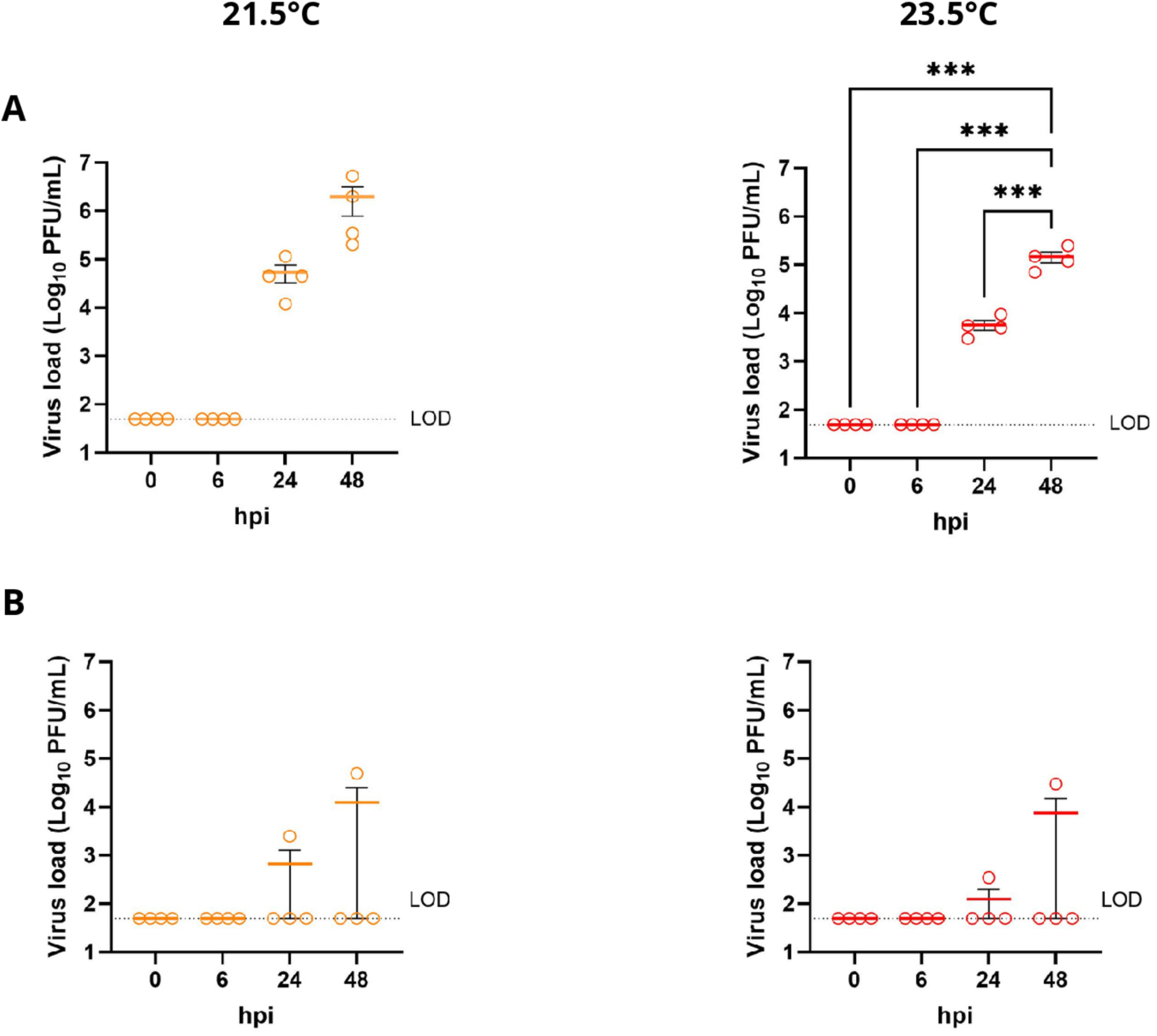
Kinetics of IHNV replication in Zf larvae at 21.5 and 23.5°C following intravenous injection or immersion infection. **A.** Infectious viral load measured at 0, 6, 24 and 48 hpi following intravenous injection at 21.5°C and 23.5°C. **B.** Infectious viral load measured at the same time points following immersion infection at 21.5°C and 23.5°C. Viral load is expressed as log₁₀ PFU/mL. Each circle represents an individual measurement (4 samples per kinetics point), and horizontal bars indicate the mean ± SEM. The dotted line indicates the limit of detection (LOD). Statistical analyses were performed using one-way ANOVA followed by Tukey’s multiple-comparisons test.

#### 3.6.4. Reverse genetic system enables the detection of the entry route of rSA-22-mCherry virus and dissemination process

Based on *in vitro* replication efficacy up to 22°C and virulence observed in Zf larvae challenged by injection and immersion at 23.5°C, the SA-22 isolate was selected for further investigation of the dynamics of the infectious process.

Given the limited proportion of infected larvae obtained following immersion challenge and the difficulty of selecting infected individuals at early stages of infection for downstream analyses (Figure 9), the SA-22 isolate was recovered using a reverse genetics approach and engineered to express a mCherry reporter cassette (Figure 10A).

**Figure 10.**
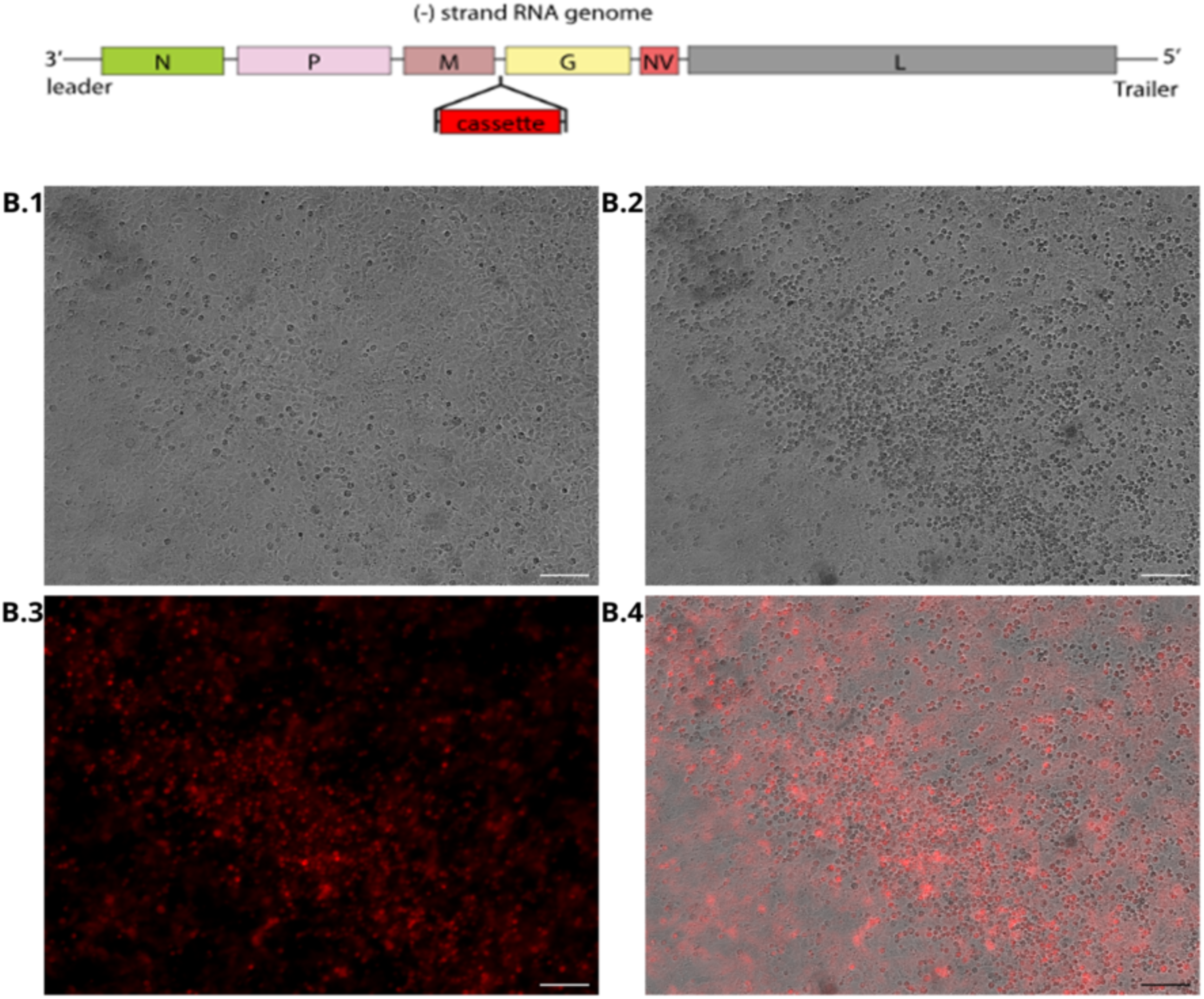
Reverse genetics system of SA-22 and detection of mCherry-positive infectious foci. **A.** Schematic representation of the rSA-22-mCherry genome engineered by reverse genetics to express the mCherry reporter gene. The mCherry expression cassette was inserted into the viral genome between the M and G genes, under the control of viral transcription signals. The recombinant virus was generated using a reverse genetics system, allowing recovery of infectious virus expressing the fluorescent reporter. **B.1.** Bright-field image showing the EPC cell monolayer of mock. **B.2.** Bright-field image showing the EPC cell monolayer at 24 hpi with rSA-22-mCherry. **B.3.** Fluorescence image of the same field as B.2 showing mCherry expression in cells infected with rSA-22-mCherry. **B.4.** Merged bright-field and fluorescence images illustrating the distribution of infected cells expressing the mCherry reporter. Images were acquired using a fluorescence microscope (5× objective). Scale bar = 100 µm.

The resulting recombinant traceable fluorescent virus enabled the direct detection and longitudinal monitoring of infected larvae following experimental challenge, providing valuable insights into viral dissemination, tissue tropism, and infection dynamics *in vivo* throughout the course of infection. The recombinant rSA-22-mCherry virus was recovered in EPC cells, reaching a final titer of 2.5 × 10⁸ PFU/mL, comparable to the wild-type (WT) SA-22 isolate. Fluorescence microscopy of EPC monolayers infected with recombinant virus revealed an intense and widespread mCherry signal, demonstrating successful reporter gene expression and confirming active viral replication. Similarly, to WT SA-22, infection with rSA-22 mCherry triggered CPE at 24 hpi at 14°C (Figure 10B). To assess the genetic stability of the recombinant virus *in vitro*, rSA-22-mCherry was serially passed twice in EPC cells. The reporter gene remained stable after these passages, with sustained fluorescence intensity, indicating maintenance of mCherry expression by the recombinant virus.

Following *in vitro* validation of the recombinant mCherry expressing virus, Zf larvae were used to investigate infection progression and viral dissemination after IV injection or waterborne immersion. The spatial distribution of rSA-22-mCherry fluorescence signal was assessed at 24 and 48 hpi by stereomicroscopy and analyzed subdividing the body axis of infected larvae into three anatomical regions: anterior, mid-body and posterior (Figure 11A). Injection of the viral inoculum into the caudal vein plexus resulted in a predominant mCherry detection in the anterior and mid-body regions at 24 hpi, in 89% and 78% of injected larvae respectively, compared with only 22% in the posterior region (Figure 11B). By 48 hpi, fluorescence was detected in all larvae across the three anatomical regions, indicating progressive systemic dissemination through the vascular network and throughout the larval body. Representative images shown in Figure 11B further demonstrated a marked increase in both the extent and intensity of fluorescence between 24 and 48 hpi, consistent with the active replication and spread of the virus.

**Figure 11.**
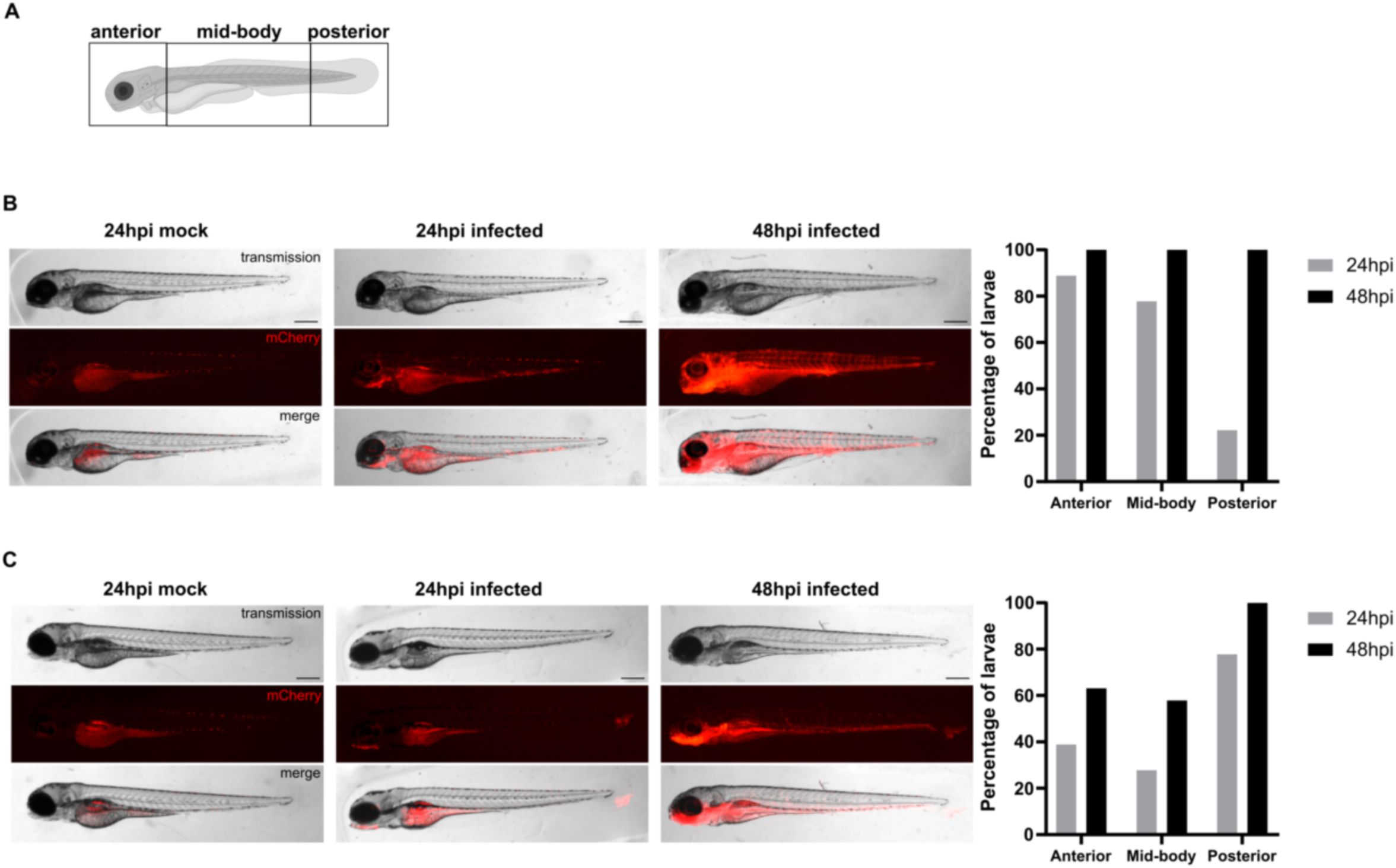
Dissemination of rSA-22-mCherry in Zf larvae at 21.5°C. **A.** Visualization of regions of interest (ROIs) for fluorescence analyses. **B.** Dissemination of rSA-22-mCherry after intravenous injection, **C.** Dissemination of rSA-22-mCherry after immersion infection. Scale bar 200μm.

In contrast, waterborne immersion resulted in a distinct distribution pattern of mCherry-positive cells detectable from 24 hpi (Figure 11C). rSA-22-mCherry fluorescence was predominantly observed in the posterior region, with 78% of positive larvae displaying mCherry-positive cells in the caudal fin. In the anterior region, mCherry-expressing cells were detected in the jaw of 39% of positive larvae. The mid-body region was less affected, with fluorescence detected in only 28% of the larvae. By 48 hpi, a marked increase in both the extent and intensity of fluorescence was observed, accompanied by an increase in the proportion of positive larvae displaying fluorescence in the anterior (63%), mid-body (57%), and posterior (100%) regions. These results indicate progressive dissemination and active replication of the virus overtime. Representative images confirmed both the increase in fluorescence intensity and the broader distribution of infected cells between 24 and 48 hpi. Overall, distinct spatiotemporal patterns of infection were observed depending on the route of administration. Direct vascular inoculation promotes rapid systemic dissemination throughout the body, whereas immersion initially resulted in localized infection of putative entry sites before progressive viral spread within the larvae.

#### 3.6.5. Caudal fin lesions were associated with IHNV infection following immersion

Macroscopic examination of Zf larvae revealed the presence of tissue lesions on the caudal fin following both intravenous injection and immersion exposure. Such lesions were observed in 42% of the rSA-22-mCherry injected larvae, compared with 45% of mock-injected larvae receiving cell culture medium (Figure 12A). In contrast, 78% of the infected larvae exposed by immersion displayed visible damages whereas no lesion was detected in the mock-infected larvae. To determine whether these tissue lesions resulted from micromanipulation during handling or from viral infection, we quantified the occurrence of mCherry-positive cells at sites of tissue damage. None of the rSA-22-mCherry injected larvae presenting caudal fin lesion (42%) exhibited detectable mCherry fluorescence within the damaged tissue. In contrast, mCherry positive cells were detected in 100 % of the larvae displaying caudal fin lesions following immersion with rSA-22-mCherry (78%). As expected, no mCherry fluorescence was observed in mock infected larvae presenting tissue damage. These results indicate that caudal fin lesions observed after immersion were strongly associated with viral infection, whereas those detected following injection were likely attributable to handling and micromanipulation procedures.

**Figure 12.**
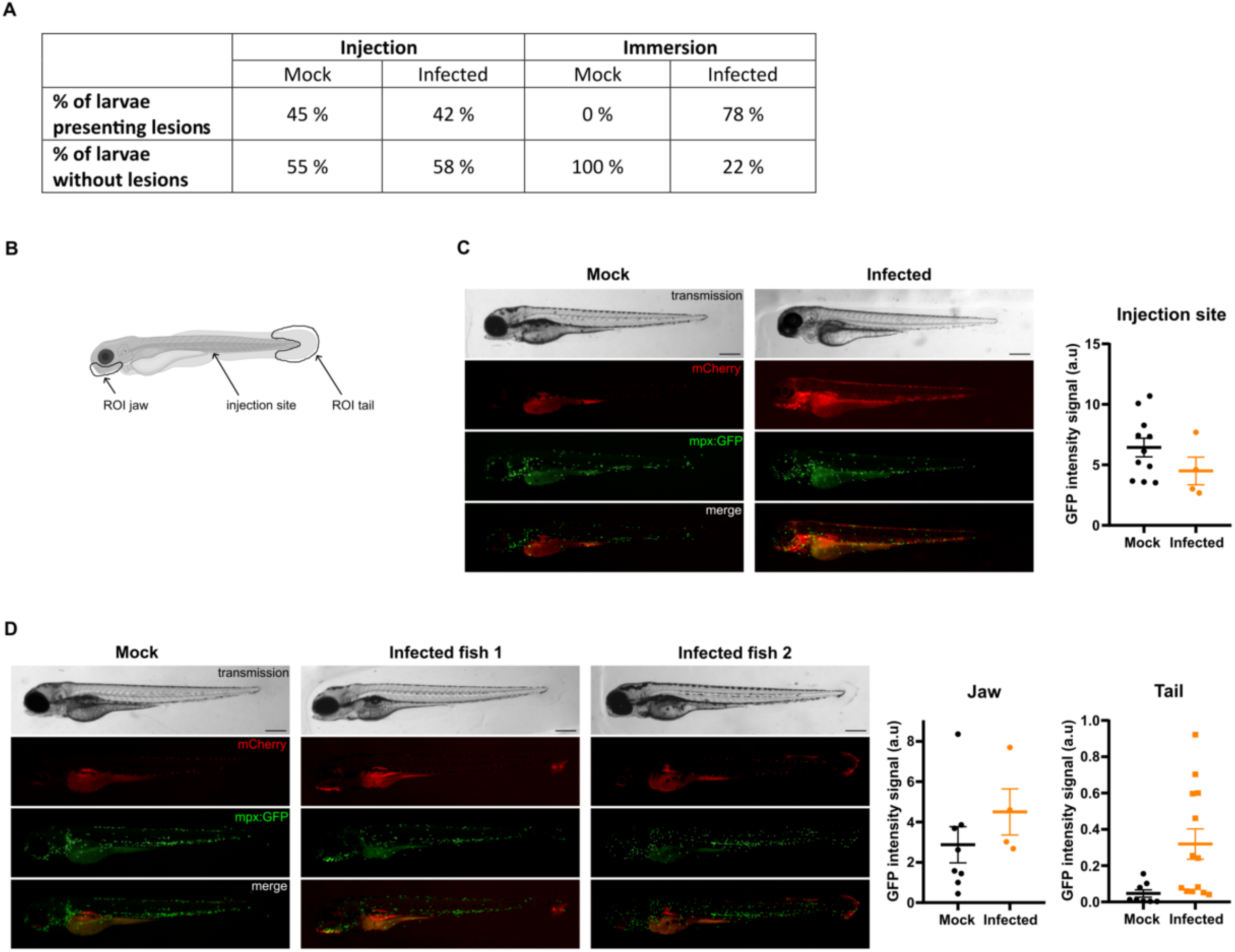
rSA-22-mCherry infection induces tissue damage and a cellular immune response in zebrafish larvae at 24 hpi and 21.5°C. **A.** Percentage of larvae presenting caudal lesion in mock-infected and infected larvae following intravenous injection or immersion. **B.** Representation of the regions of interest used for GFP fluorescence-intensity quantification, including the intravenous injection site and the two primoreplication sites identified following immersion exposure to rSA-22-mCherry, the jaw and the caudal fin. **C.** Representative images of rSA-22-mCherry dissemination following intravenous injection, showing bright-field, mCherry fluorescence, *mpx*:GFP fluorescence and merged images. Quantification of GFP fluorescence intensity at the injection site is shown on the right. **D.** Representative images of rSA-22-mCherry dissemination following immersion infection, with bright-field, mCherry, *mpx*:GFP and merged images. Quantification of fluorescence intensity in the jaw and tail is shown on the right. Data are presented as individual values with mean ± SEM. Scale bars, 100 µm. Mann–Whitney test: non-significant difference.

#### 3.6.6. In contrast to intravenous injection, waterborne infection route promotes neutrophil recruitment to the infectious foci

As tissue damage is known to trigger neutrophil recruitment as part of the wound-healing response, we next sought to characterize the cellular response at the mCherry positive infection site. To this end, *Tg(mpx:GFP)* Zf larvae, in which neutrophils express GFP, were infected with rSA-22-mCherry. In immersed larvae, neutrophils were consistently recruited to the caudal-fin lesion in all mCherry-positive individuals.

As expected, mock infected larvae showed no GFP fluorescence in the caudal fin in absence of tissue lesion. For injected larvae, caudal fin lesions occurred at similar frequencies in infected and mock-injected controls (Figure 12A), but no mCherry signal was detected at this site. Nevertheless, GFP-positive neutrophils were present at the lesion in 37 % of infected larvae, whereas mock-injected larvae showed no GFP signal. Together, these observations indicate that immersion triggers a consistent, lesion-associated recruitment of GFP-positive neutrophils linked to mCherry-positive infection, whereas lesions observed in injected larvae occur independently of infection status but elicit GFP-positive neutrophils in only a subset of infected larvae.

Neutrophil recruitment was next quantified at 24 hpi by designating three regions of interest (ROIs): the intravenous injection site (caudal venous plexus), and the two sites of primo-replication of the virus (the caudal fin and the jaw) (Figure 12B), which became mCherry-positive as early as 24 hpi following immersion infection. At this stage, GFP signal intensity at the intravenous injection site was not significantly different between larvae injected with rSA-22-mCherry and those injected with culture medium, suggesting that the presence of the virus does not induce local cellular response following injection (Figure 12C). In larvae infected by immersion, GFP fluorescence intensity was higher at the mCherry-positive primo-replication sites than in the corresponding regions examined in the mock-infected larvae, although the difference did not reach statistical significance (Figure 12D).

Together, these results suggest that the host immune response is shaped by the route of viral entry. Waterborne immersion induced a local leukocyte recruitment at putative sites of viral entry, particularly in association with infected tissues, whereas this response was not observed following systemic inoculation. This indicates that mucosal or peripheral exposure to IHNV elicits an early localized immune response that differs from that triggered by direct viral delivery into the circulation.

## Discussion

IHN is a major EU-regulated disease of salmonids, whose clinical expression generally occurs at water temperatures below 14°C (Amend, 1970; Hbtrick et al., 1979; Mulcahy et al., 1984). The etiological virus appears to have a faster evolution potential than other salmonid viruses such as the related Viral Hemorrhagic Septicemia Virus (VHSV) and Infectious Pancreatic Necrosis Virus (IPNV) (Abbadi et al., 2016; Cieslak et al., 2017). In recent years, an increase in IHNV incidence, together with the spread of the infection into previously disease-free northern European countries has been reported in parallel with the growing impacts of global warming (Abaddi et al., 2021; Kantala et al., 2026). To contribute to the evaluation of the possible effects of rising water temperature on the epidemiology of this regulated disease, we investigated in this work the replication thermoprofiles of six IHNV field isolates selected based on their genotype (M, L, E and J), geographical origin, years of isolation and host species, using a combination of *in vitro* and *in vivo* approaches. Distinct thermoprofiles were identified across the 4 evaluated genotypes based on their ability to replicate *in vitro* at 14°C and warmer sub-optimal temperatures (20, 22 and 24°C). Pre-screening based on the total viral production combining intracellular and extracellular infectious particles revealed thermo-sensitivity at temperatures of 20°C and above for two isolates (FR0031 genotype L and 1075b, E) whereas all others retained detectable replication at this temperature. None of the isolates replicated significantly at 24°C, indicating that this temperature likely represents the upper thermal limit for productive IHNV replication in EPC cells, a range consistent with recent observations by Marsella et al. (2025). Based on these results, we conducted more detailed analyses on SA-22 (genotype E, isolated in 2022) and RtNag0a (genotype J, isolated in 2006) - two isolates selected for their replication competence at 20 but also 22°C along with N61 (E), the French reference strain isolated in 1989. Pairwise comparison of the complete genome sequences revealed a high level of genetic similarity among the three IHNV isolates (nucleotide and amino acid identities of 96.35% and 96.33% respectively between N61 and SA-22; 95.74% and 95.73% between N61 and RtNag0a; 93.54% and 93.47% between SA-22 and RtNag0a), indicating a closely relation at the whole-genome level, while showing a degree of genetic divergence consistent with their distinct evolutionary backgrounds.

We thus quantified intracellular and extracellular viral RNA released in the EPC supernatant (containing infectious virions but also defective viral particles) during the first 72 hours following infection, thereby characterizing the temperature-dependent replication profiles of these isolates. SA-22 consistently exhibited the highest levels of extracellular viral RNA particularly at 20 and 22°C, whereas N61 was more strongly affected by the increase in temperature. RtNag0a exhibited an intermediate phenotype. Additionally, strain SA-22 exhibited the lowest ratios of infectious particles to genome copies in the extracellular medium, regardless of temperature or kinetic time point (data not shown). In contrast, intracellular viral RNA accumulation was particularly pronounced for SA-22 but also RtNag0a at the highest temperatures. This discrepancy between intracellular and extracellular yields and between infectious particles and genome copies ratios for extracellular compartment, implies that viral genetics and temperature may influence one or more stages of the viral cycle. Although the specific mechanisms cannot be resolved from current results, temperature-sensitive factors such as RNA structural dynamics, protein–RNA and protein–protein interactions, or changes in the conformation and activity of viral enzymes, could in turn influence viral genome replication and downstream processes of the viral cycle, including genome packaging, particle assembly or release (Bisht & Velthuis, 2022).

Temperature-dependent susceptibility to infection is also shaped by the cellular environment, an aspect addressed in our study using both fathead minnow EPC cells, and rainbow trout–derived RTG-2 cells. These two lines provide distinct physiological contexts, which may differentially modulate viral replication dynamics even though both support rhabdovirus infection. Our data further demonstrate the replicative capacities of thermotolerant isolates at 14 and 20°C in RTG-2, showing that these trout-derived cells provide a more permissive environment for viral replication at 20°C, with replication rates exceeding those observed at 14°C and surpassing those measured in EPC cells at 20°C. These observations suggest that the physiological features intrinsic to cellular models modulate viral replication efficiency, thereby contributing to the differential thermal sensitivity observed across isolates. Moreover, systematic assessment of viral replication across multiple cell lines including EPC and RTG-2 is essential to capture host specific permissiveness and to better understand how cell environment diversity shapes viral performance under varying thermal conditions.

Unexpectedly, the replicative profiles of the isolates were not directly reflected by the severity of CPE in EPC cells, which varied according to the temperature and the time post infection. These observations likely reflect an interplay between isolate-specific virulence and the monolayer’s healing capacity, both influenced by temperature (Pumputis et al., 2022). Indeed, temperature can also modulate EPC cell physiology, including metabolic fluxes, membrane behavior, intracellular transport systems and innate immune competence. Our study examined key cellular stress and antiviral pathways, including interferon signaling through *isg15* expression (Langevin et al., 2013), oxidative stress response mediated by *nrf2* (Yang et al., 2014) and endoplasmic reticulum stress monitored via *atf6* (Salvador-Mira., 2024). Temperature exerted a clear influence on both the magnitude and the kinetics of the cellular response to IHNV infection, particularly for *isg15* and *atf6*. At 14°C, *isg15* showed only modest activation, primarily at 24 hpi, whereas markedly stronger and earlier induction occurred at higher temperatures, with responses shifting from 72 hpi at 14°C and 20°C to as early as 48 hpi at 22°C. A similar temperature-dependent acceleration was observed for *atf6,* with a gradual induction between 24 and 48 hpi at 14 °C, in contrast to a strongly elevated and sustained induction detected at 20°C and 22°C from 24 hpi onward. In contrast, *nrf2* displayed a progressive but largely temperature-independent pattern of activation, increasing from 24 hpi and remaining elevated at 48 hpi under all conditions. Together, these results indicate that while higher temperatures accelerate and amplify antiviral and ER-stress responses, oxidative-stress signaling through *nrf2* follows a more stable temporal trajectory, suggesting that distinct cellular pathways differ in their sensitivity to thermal modulation during IHNV infection. However, the expression profiles of these markers did not necessarily follow the same kinetics as viral production, highlighting the complex interaction between viral replication, temperature and host cell responses.

These observations are in agreement with recent work examining the influence of the water temperature on IHNV replication and intestinal immunity in rainbow trout (Hai et al., 2026). In that work, IHNV replicated more efficiently at 12.5°C than at 16.5°C, yet mucosal immune enzyme activities and intestinal transcriptomic profiles showed marked temperature-dependent differences. The intestinal transcriptional response was more extensive at 12.5°C, with temperature-specific differentially expressed genes enriched in the PI3K–Akt signaling pathway, suggesting that IHNV infection, optimal at lower temperatures, exerts a stronger and potentially dysregulated impact on host immunity under cold conditions. Enzymatic activities such as ACP, AKP, LZM, and SOD also varied with temperature, with fish at 12.5°C exhibiting stronger suppression of hydrolytic immune functions and higher oxidative-stress responses than those at 16.5°C. Together, these findings indicate that lower environmental temperatures can exacerbate the physiological impact of IHNV infection, potentially enhancing viral pathogenicity or facilitating immune evasion while simultaneously driving a more intense but less effective host response.

A similar temperature-dependent modulation of viral infection and immunity has been described for VHSV, where viral replication and innate responses in primary flounder spleen cells varied across temperatures (Jang et al., 2025). Although *in vivo* phenotyping identifies 15°C as the optimal replication temperature and 25°C as non-permissive, VHSV replication can be observed *in vitro* particularly at 15°C and 20°C. Viral loads differed by more than ten-fold for each 5°C increment, suggesting that reduced activity of the viral RNA-dependent RNA polymerase at higher temperatures may limit replication despite ongoing infection. Innate immune responses also showed temperature-dependent modulation, pro-inflammatory cytokines peaked at 15°C, *isg15* and *mx* expression were slightly higher at 20°C, and both cytokine and interferon-related gene expression were markedly lower at 25°C despite detectable viral replication. These VHSV data parallel our IHNV observations and illustrate that temperature can uncouple viral replication from the strength and timing of host-cell responses, reinforcing the idea that thermal conditions shape not only viral performance but also the quality, magnitude, and coordination of antiviral pathways.

Taken together, our study enabled the characterization of field isolates that retain replication capacity *in vitro* at temperatures classically considered non-permissive, supporting experimental data and recent surveillance reports indicating that IHNV can circulate or cause mortality beyond its canonical temperature range (Vendramin et al., 2022; Marsella et al., 2025). While studies conducted as early as the 1970s examined the *in vitro* thermal replication profiles of North American IHNV isolates, demonstrating replication at 20°C but not at 23°C (Wingfield et al., 1969; Mulcahy et al., 1984), the first dedicated investigation of European genotype E isolates was only recently published by Marsella et al. (2025). Among the six European isolates examined in their study, two showed marked thermotolerance, namely IHNV DK, isolated in 2021 during surveillance in Denmark at a water temperature of 19°C, and IHNV 67-1, isolated in 2022 during an outbreak in Italy at a water temperature of 16-18°C. Both isolates were capable of replicating at 22°C in EPC, highlighting their elevated thermal tolerance relative to the canonical IHNV replication range. Our results extend the panel of thermotolerant IHNV isolates by identifying the French genotype E isolate SA-22, isolated in 2022, as well as the Japanese genotype J isolate RtNag0a, both of which retained viral replication at elevated temperatures.

The phylogenetic analysis based on complete G-gene nucleotide sequences further showed that SA-22 clustered within the monophyletic group formed by IHNV 67-1, IHNV DK, and IHNV 217/A, all of which replicate at elevated temperatures *in vitro*. This positioning reinforces the hypothesis of a shared evolutionary background associated with thermotolerance proposed by Marsella et al. (2025). In contrast, RtNag0a was positioned separately within genotype J. Moreover, a distinct seven amino acids insertion in the 3′ end of the L gene was identified in both IHNV 67–1 and IHNV DK isolates, potentially representing a molecular signature linked to enhanced replication capability and adaptation to suboptimal temperatures (Marsella et al., 2025). This extension was absent from SA-22 and RtNag0a, even though both isolates replicate at elevated temperatures. However, SA-22 exhibited alteration within the L-protein C-terminus, notably a premature stop codon shortened the L protein by one amino acid, which was not found in the thermosensitive European isolates (Italian and French), the Japanese RtNag0a isolate, or the IHNV25 *in vitro* thermo-adapted variant. This divergence reinforces the hypothesis that adaptive structural remodeling of the L-protein C-terminal domain might play a pivotal role in thermotolerance. It would be particularly interesting to determine, using reverse genetics approaches, the involvement of these mutations in the thermotolerance phenotype of this isolate. The in *vitro* thermo-adapted IHNV25 also displayed several amino acid substitutions in the L protein, without sharing the C-terminal extension, indicating that there is no single mutational pathway to obtain thermotolerant strains. While the L protein has been shown to influence temperature-dependent growth in *Novirhabdovirus* (Kim et al., 2015), the G protein has also been investigated as a potential determinant of the thermal restriction of IHNV. However, Biacchesi et al. (2002) showed that replacement of the IHNV *G* gene with the SVCV G gene, from a virus capable of replicating at temperatures up to 28°C, did not expand the permissive temperature range of the recombinant IHNV. Overall, the genetic comparison of field and *in vitro* thermo-adapted isolates revealed distinct sequence patterns associated with elevated-temperature replication. A comprehensive genomic comparison of isolates displaying contrasting thermoprofiles, combined with reverse genetics approaches, will therefore be required to identify the viral genes involved as well as the molecular determinants underlying IHNV thermotolerance.

Consistent with findings reported for VHSV (Jang et al., 2025, Marsella et al. (2025) also highlighted that *in vitro* thermoprofiles established in EPC do not necessarily predict *in vivo* virulence in rainbow trout challenged at different temperatures. Their experimental challenges showed that IHNV 67-1 induced mortality at both 12 and 19°C, whereas IHNV DK induced mortality only at 12°C, illustrating the complexity of the relationship between viral thermal phenotype and disease outcome in the host.

In this context, we examined the virulence of N61, SA-22, and RtNag0a selected based on their *in vitro* thermoprofiles, to determine whether thermal tolerance in cell culture translates into pathogenicity *in vivo.* In compliance with animal experimentation legislation and in strict application of the 3Rs principles (Replace, Reduce, Refine), IHNV virulence was investigated in Zf larvae, a validated model for experimental infections with both IHNV and VHSV (Aggad et al., 2009; Ludwig et al., 2011; Marana et al., 2021). Zf were reared at 28°C under standard conditions, but the temperature was reduced to 24°C for IHNV to establish robust infectious models supported by a thermo-adapted variant IHNV 25-70. Despite the developmental delay caused by lowering the temperature, Zf larvae retain the ability to elicit antiviral responses, notably through interferon signaling and neutrophil activation. This immunocompetence allows the investigation of virulence and host– pathogen interactions in physiological conditions. Intravenous inoculation of thermo-adapted IHNV variants (IHNV 25-70 and IHNV25) generated *in vitro* through iterative passages up to 25°C, led to efficient replication in larvae at 24-25°C and induced clinical signs culminating in the death of the infected larvae within 3–4 dpi (Ludwig et al., 2011; Fauvet et al. submitted).

Interestingly, the impact of temperature on IHNV pathogenesis further supports the importance of early events during infection. A temperature shift performed at 6 hpi from 24°C to 28°C markedly reduced viral dissemination and was sufficient to prevent mortality, suggesting that establishment of IHNV infection relies on early mechanisms of viral replication and immune evasion, highly sensitive to environmental conditions (Ludwig et al., 2011). Therefore, our understanding of IHNV infection dynamics and host antiviral responses is predominantly derived from Zf models using laboratory thermo-adapted viral strains, capable of replicating at temperatures above 20°C. Consequently, the extent to which these findings accurately reflect the biology and host interactions of circulating wild-type isolates remains to be established. In the present study, larvae were successfully infected with field isolates SA-22 and RtNag0a and the reference strain N61 without the need for prior thermal adaptation. The thermotolerant profiles of these isolates in larvae, where SA-22 and RtNag0a have been replicated with similar efficiency at 21.5 and 23.5°C whereas replication of the thermosensitive reference strain N61 was impaired at 23.5°C, are consistent with those observed in EPC. Following intravenous infection, the thermotolerant isolates SA-22 and RtNag0a induced mortality kinetics similar to those previously reported for IHNV 25-70 (Ludwig et al., 2011), reaching approximately 90% cumulative mortality within 3–4 days. SA-22 induced higher cumulative mortality at 23.5°C than at 21.5°C at 48 hpi, although this difference became less pronounced by 72 hpi. In contrast, RtNag0a tended to cause more rapid mortality at 21.5°C than at 23.5°C at 48 hpi, although the difference was not statistically significant, and ultimately reached similarly high cumulative mortality rates at both temperatures. In contrast, the thermosensitive reference strain N61 exhibited delayed mortality kinetics, with only approximately 50% cumulative mortality at 96 hpi. Macroscopic examination of infected larvae enabled the monitoring of clinical signs onset and confirmed the pathophysiological progression of the disease as previously described by Ludwig et al (2021). Edema and hemorrhages were observed at 21.5°C but were even more pronounced at 23.5°C, with the appearance of exophthalmos, a hallmark clinical sign typically reported in IHNV-infected salmonids. This increased disease severity may reflect more rapid viral dissemination or a more deleterious host response at higher temperature, despite comparable survival times.

Similarly, to IHNV 25-70 injected intravenously, the field isolates SA-22 and RtNag0a elicited a clear activation of anti-viral IFN signaling pathways, as evidenced by the induction of *ifnφ1, ifnφ3* and IFN-induced genes *viperin/rsad2* at 48 hpi, together with a robust upregulation of the pro-inflammatory cytokine *il1b* gene. Ludwig et al. (2011) concluded that this delayed interferon activation supports the existence of IHNV-mediated anti-IFN mechanisms that dampen the host response during the early stages of infection (Ludwig et al., 2011). Consistent with this hypothesis, Aggad et al. demonstrated that administration of recombinant Zf IFNφ1 and IFNφ2 to larvae prior to IHNV challenge increased survival in a dose-dependent manner, indicating that an early IFN response is sufficient to efficiently restrict viral infection (Aggad et al., 2009). Notably, the similar pattern of IFN pathway activation observed for both the thermo-adapted IHNV strain and the non-adapted yet thermotolerant field isolates suggests that thermotolerance of IHNV, whether experimentally acquired or naturally occurring, is accompanied by a conserved host response architecture, reinforcing the notion that temperature associated viral fitness primarily influences infection efficiency and modulates both the magnitude and the kinetic of the antiviral response.

In addition to the unexpected ability of these field isolates to replicate *in vivo* at 23.5 °C, a temperature previously believed to support infection only by the thermo-adapted IHNV 25-70 and IHNV25 variants, waterborne exposure to SA-22, RtNag0a, and, to a lesser extent, N61, resulted in clinical disease and mortality, although less efficiently than intravenous inoculation and affecting a smaller proportion of individuals. Nevertheless, these findings challenge earlier conclusions that immersion is an ineffective route of IHNV infection in Zf larvae and instead demonstrate that non-adapted yet thermotolerant isolates can efficiently initiate infection under conditions previously considered restrictive (Ludwig et al., 2011).

Several factors may explain these differences in susceptibility to waterborne models: genetic and virulence of IHNV isolates, impact of water temperature on Zf immune competences and/or composition of the fish microbiota. The influence of temperature on innate immune response has been addressed in several studies. Zhang et al., for instance, have investigated the inflammatory and immune responses induced by lipopolysaccharide (LPS) on early larval stages maintained at different thermal regimes (24, 28, or 32 °C) from embryogenesis to the onset of first feeding (Zhang et al., 2018). Survival following LPS challenge strongly depended on the embryonic incubation temperature, 24°C displaying the highest mortality rate compared to 28°C or 32°C, regardless of the temperature at challenge. Transcriptomic analyses further indicated that low embryonic temperatures impair the effectiveness of innate immune defenses despite inducing inflammatory responses, whereas higher incubation temperatures promote a stronger immune activation. Conversely, the highest challenge temperature dampened immune responses and stimulated hypoxia and oxidative stress related pathways. These observations highlighted the strong influence of temperature on the Zf innate immune competence and suggest that the susceptibility of larvae to IHNV infection may reflect not only viral thermotolerance but also that temperature experienced during embryogenesis (28°C) can shape host susceptibility to subsequent pathogen challenge at 21.5 and 23.5°C. Interestingly, thermal choice experiments have provided further evidence of the influence of temperature on Zf immune responses. Using a water-column system with a vertical temperature gradient ranging from 28°C to 32°C, Zf larvae showed little thermal discrimination whereas later stages of developments (>13 dpf) preferred temperatures of 30–31°C (Rey et al., 2017). Following challenge with double-stranded RNA (dsRNA), which mimics replication intermediates during RNA virus infections, fish displayed a late-onset behavioral fever response migrating to warmer waters (32°C), only at 18–20 dpf. Importantly, these fish exhibited stronger induction of antiviral genes than challenged fish maintained at a constant temperature of 28°C, supporting previous results demonstrating that temperature is a major regulator of teleost immune competence, influencing both innate and adaptive immune functions, pathogen susceptibility, and the capacity of the host to mount effective antimicrobial and antiviral responses (Rakus et al., 2017; Scharsack & Franke, 2022).

Beyond its effects on molecular and transcriptional immune responses, temperature strongly influences the recruitment, migration, and effector functions of innate immune cells (Metz et al., 2026). Using the Zf tail fin amputation model, Metz et al. showed that temperatures between 22 and 35°C substantially modulate neutrophil and macrophage dynamics. Neutrophils recruitment increased progressively with temperature, whereas macrophage recruitment displayed a thermal optimum around 31°C, which also corresponded to maximal tissue regeneration. In contrast, higher temperatures delayed inflammation resolution, promoted persistent neutrophil accumulation, and impaired tissue repair. This dynamic study demonstrates that temperature is a key regulator of innate immune cell behavior and, consequently, of host responses to infection. In addition to its effects on immune cell behaviors, temperature can also indirectly modulate disease susceptibility by shaping the composition and activity of the host microbiota (Zhang et al., 2024). In Zf, high thermal condition increased the abundance of *Parabacteroides distasonis* and promoted the production of deoxycholic acid (DCA), a metabolite that suppresses the infectivity and pathogenicity of SVCV, another fish rhabdovirus. Importantly, DCA was subsequently identified as a potent inhibitor of IHNV infection in rainbow trout, highlighting a previously unrecognized link between temperature, composition of gut microbiota and antiviral resistance mechanisms in fish. Together, these findings highlight the multifaceted and highly context-dependent effects of temperature on host–pathogen interactions. Temperature influences not only viral replication but also immune competence, immune cell dynamics, and host–microbiota interactions, all of which contribute to determining disease outcome. Consequently, the impact of temperature on resistance and susceptibility to viral infections is complex and multifactorial. Further studies are therefore needed to better understand the mechanisms through which temperature shapes antiviral defenses and disease susceptibility, particularly for viral pathogens responsible for significant economic losses in aquaculture.

Recovery of rSA-22-mCherry opened up opportunities to decipher the dynamic of the infectious process of this isolate of particular interest *in vivo* using fluorescence imaging of Zf larvae. The identification of initial-replication sites within the superficial tissues of the caudal fin and jaw is fully coherent with the established entry sites described for IHNV in salmonids exposed to the virus by the natural waterborne route (Harmache et al., 2006). In these species, IHNV targets several key portals of entry including the skin, fin bases, and the oral region/esophagus–cardiac stomach region, where initial replication occurs in epidermal cells before the virus disseminates to hematopoietic tissues, blood capillary endothelial cells, the kidney, and other internal organs (Harmache et al., 2006; Schönherz et al. 2012; Tulaby Dezfuly et al., 2026). Our model therefore enables the observation of early infection phases initiated at peripheral epithelial barriers, consistent with the tissue tropism revealed by recombinant IHNV-luciferase imaging in salmonids (Harmache et al., 2006). Virus dissemination was readily visualized, and contrasted phenotypes were observed between exposure routes. Zf challenged by intravenous injection exhibited rapid, systemic spread, whereas fish infected by immersion displayed a distinct dissemination pattern, with infection progressing from peripheral tissues. This divergence in kinetics and tissue tropism highlights the fundamentally different early infection dynamics associated with intravenous inoculation versus waterborne exposure. Furthermore, fluorescent transgenic lines offered a powerful means to visualize neutrophil behavior *in vivo*, enabling real-time monitoring of their recruitment, migration, and tissue-specific activity. These models thereby facilitated detailed analysis of immune activation triggered by immersion challenge, in which neutrophils are rapidly recruited to infectious foci within superficial epithelial layers, a pattern that differs markedly from the response observed at the injection site. This contrasted spatial pattern of neutrophil recruitment thus highlights the distinct immune responses elicited depending on the route of entry, with waterborne infection engaging peripheral mucosal defenses and systemic infection triggering a different inflammatory architecture. Together, these observations support a more integrated understanding of early host–pathogen interactions and the spatial dynamics of neutrophil recruitment and mucosal immunity during IHNV infection (Marana et al., 2021; Souto et al., 2024). In light of our findings demonstrating the thermotolerance of the N61, SA-22 and RtNag0a isolates, it will be important to re-evaluate and complete previous studies characterizing the course of infection and host immune responses in rainbow trout, conducted with N61 and RtNag0a at 11°C (Louboutin et al., 2021). Future challenge experiments performed at higher temperatures, including 20°C, will be necessary to assess the infectious risk posed by these field isolates in warm conditions. Such studies should determine whether increased temperature alters virulence, symptomatology and mortality rates, hematological parameters, and the magnitude and kinetics of interferon-mediated antiviral responses and pro-inflammatory pathways. More broadly, revisiting the pathogenicity of these isolates across a wider thermal range will provide a better understanding of their epidemiological potential under current aquaculture conditions and in the context of ongoing climate warming.

In conclusion, we showed that some IHNV field isolates exhibit marked specific replication thermoprofile influenced by the viral genetic background and the cellular environment. While all isolates replicated efficiently at 14–20°C, their capacity to maintain productive replication at higher temperatures revealed thermotolerant (SA-22 and RtNag0a) and thermosensitive (N61) isolates in EPC cell model. Modulation of these observations in RTG-2 cells highlights the importance of host cell physiology in viral fitness. Furthermore, we showed that replication efficiency and cytopathogenicity are not necessarily coupled, supporting the existence of distinct thermal adaptation mechanism among IHNV isolates. Such phenotypic diversity emphasizes the importance of considering both viral and host factors when assessing the epidemiological risk posed by emerging IHNV variants in aquaculture. Zf proved to be an elegant and efficient *in vivo* model to assess the virulence of our naturally thermotolerant isolates at 21.5 and 23.5°C, with mortality kinetics correlating with replication levels observed *in vitro* on EPC cells. This *in vivo* model proved to be efficient for infection by immersion using field viral isolates for the first time – without requiring additional *in vitro* thermo-adaptation - and further allowed the study of viral spreading and neutrophil mobilization by imaging. Altogether, these results enhance our understanding of how rising environmental temperatures may drive the emergence of thermotolerant IHNV variants, thereby improving our ability to anticipate epidemic risks under future climate scenarios. Experimental infection trials incorporating the effects of temperature on fish physiology and immune response are in progress to complete the risk assessment of these thermotolerant isolates on a major IHNV target species - rainbow trout.

## Acknowledgments

We would like to thank Evelyn EMMENEGGER and Gael KURATH, from the US Dept. of Interior, US Geological Survey Western Fisheries Research Center, Seattle, USA, who kindly agreed to provide us a set of 24 strains of IHNV, among which three were used in the study described here. We also thank the European Reference Laboratory for Fish and Crustacean Diseases, DTU-aqua Denmark, who kindly provided us with the complete nucleotide sequence of the IHNV DK-FF1/21–4070–1 isolate. We are grateful to members of the fish facilities, Magali Bouvet, Pénélope Simon, Damien Degallaix and Kilian Cella, for taking care of experimental fish (IERP-UE907, Jouy-en-Josas Research Center, France doi.org/10.15454/1.5572427140471238E12) which belongs to the National Distributed Research Infrastructure for the Control of Animal and Zoonotic Emerging Infectious Diseases through In Vivo Investigation (EMERG’IN DOI: doi.org/10.15454/90CK-Y371).

## Author Contributions

Author contributions following the CRediT taxonomy (https://casrai.org/credit/) are as follows: Conceptualization: CL, TM, SB; Formal analysis: CL, TM, EG; Funding acquisition: CL, TM, SB, DR, YB; Investigation: EG, CF, MM, PP, MA, MF, SB, JKM, LL, GL, YB, MLT, PF, DR, TM, CL; Methodology: EG, CF, MM, PP, MA, MF, LL, GL, MLT, PF; Project administration: CL, DR, TM, JKM, SB, YB; Supervision: CL, TM, SB, JKM; Visualization: EG, CL, MM, MA, LL, TM; Writing – original draft: CL, TM, EG; Writing – review & editing:. EG, CF, MM, PP, MA, MF, SB, JKM, LL, GL, YB, MLT, PF, DR, TM, CL

## Data availability

Data will be made available on request.

